# Reversing PROTAC-induced ASH2L degradation reactivates proliferation in senescent cells

**DOI:** 10.64898/2026.05.02.722411

**Authors:** Janina Müller, Mirna Barsoum, Philip Bussmann, Mohamed H. E. Mabrouk, Roksaneh Sayadi, Anna-Maria Vllaho, Alexander T. Stenzel, Lukas Vogt, Paul Kiessling, Christoph Kuppe, Juliane Lüscher-Firzlaff, Bernhard Lüscher

## Abstract

Nucleosomes control access to gene promoters. Histone H3 lysine 4 tri-methylation, catalyzed by 6 KMT2 complexes, correlates with accessible promoters and gene expression. The catalytic activity of KMT2 enzymes depends on an obligatory core complex with ASH2L being an essential subunit. We find that PROTAC induced depletion of ASH2L reduces H3K4me3, deregulates gene expression and prevents proliferation. Upon prolonged ASH2L loss, cells develop a senescent phenotype, a process linked to aging and disease. Competing the PROTAC reactivates ASH2L, reestablishes H3K4me3 at promoters and reverts gene expression changes. Cells reenter the cell cycle and resume proliferation, thereby reverting senescence. Structure-function studies demonstrate that these molecular and cellular consequences are primarily due to the loss of ASH2L functions associated with KMT2 complexes. Together, these findings indicate that stress inflicted by the loss of KMT2 catalytic activities promotes a reversible senescence phenotype, suggesting that the functions of KMT2 complexes are implicated in aging.

**Graphical abstract:** 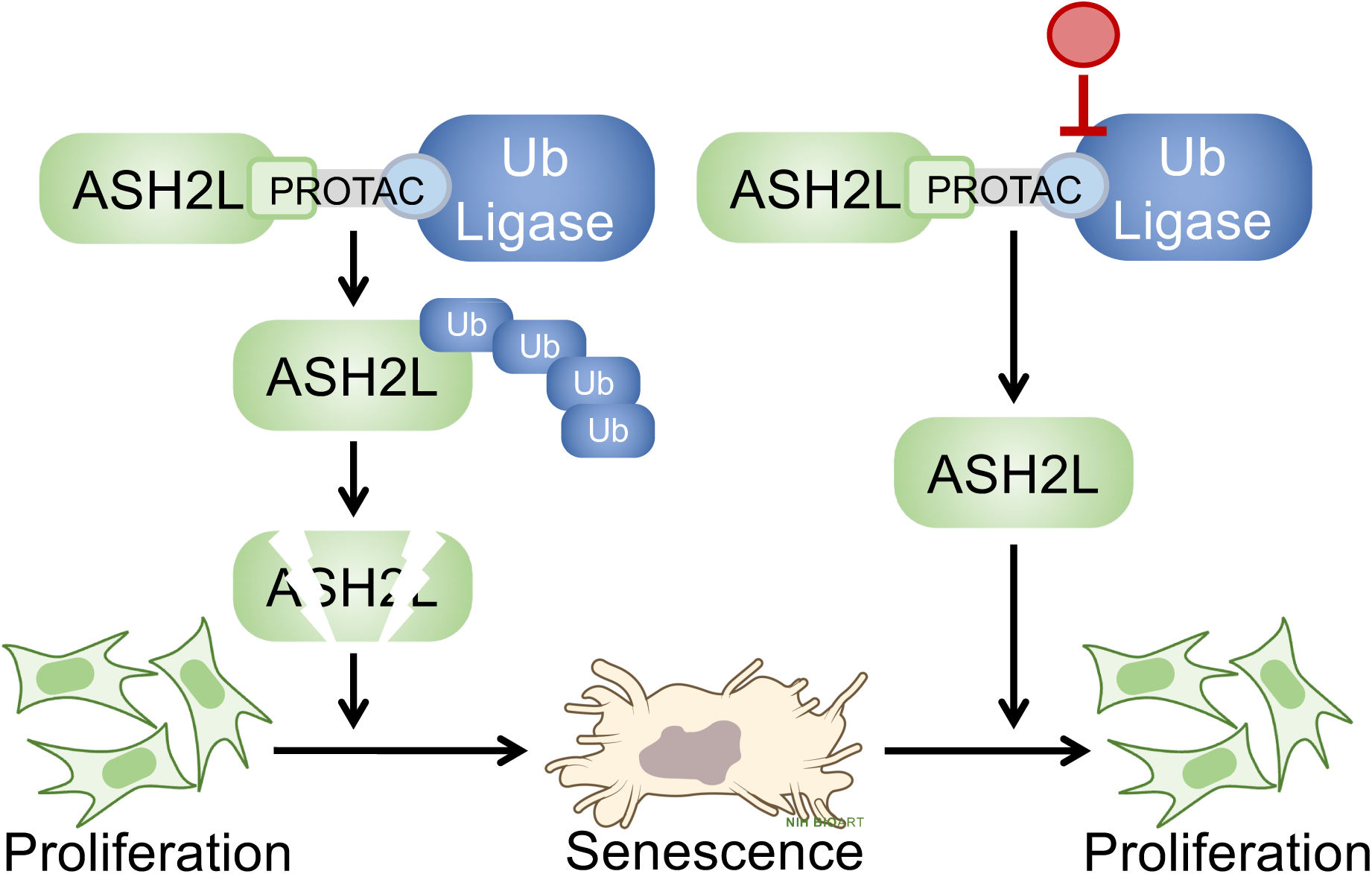

## Introduction

Cell development and identity is dependent on differential regulation of gene expression, which is controlled by transcription factors and by chromatin, the smallest unit being the nucleosome (1–5). Nucleosomes restrict the accessibility to DNA and thus have gate-keeper function for controlling DNA-associated processes, such as transcription (6). Nucleosomes with their histone protein core function as platforms for factors that affect transcription. These interactions are regulated by a multitude of post-translational modifications on core histones (PTMs), whose functions are only partially understood, particularly when combinatorial effects are considered (5,7–9).

Certain histone PTMs, also referred to as histone marks, are linked to open, potentially transcribed chromatin, while others are found in more compact heterochromatin. For example, in the latter tri-methylation of histone H3 lysine 9, lysine 27 and histone H4 lysine 20 (H3K9me3, H3K27me3, and H4K20me3, respectively) are found, along with the presence of the linker histone H1 (10). Chromatin regions with active transcription contain typically acetylation marks of multiple lysines of different core histones and methylation of H3K4 (7). The latter has gained considerable attention as H3K4me1 is associated with active enhancers and H3K4me3 with active promoters, establishing a strong correlation of H3K4 methylation with gene transcription (11–15). This suggests that the enzymes that catalyze (writers) and remove (erasers) these PTMs as well as the readers, which disseminate the information associated with H3K4 methylation, are important regulators of gene transcription.

The H3K4 methyltransferases associated primarily with transcription belong to the KMT2 family, which consists of 6 catalytically active members. All 6 enzymes associate with the WRAD core complex, composed of WDR5, RBBP5, ASH2L and DPY30, which is essential for efficient catalytic activity (11–15). The close association with transcription has been supported by studies of individual KMT2 enzymes, while more global analyses have been achieved by manipulating WRAD subunits. The knock-out (KO) of *Ash2l* or *Dpy30* has broad effects, preventing cell proliferation and differentiation, and both are necessary for tissue integrity (16–21). However, due to the stability of the encoded proteins, the effects develop slowly and, therefore, direct mechanistic studies are not straightforward.

The recent development of PROTAC (proteolysis targeting chimeras) approaches to deplete RBBP5, ASH2L and DPY30 has advanced the molecular understanding of KMT2 or COMPASS-like complexes (22–24). All three proteins are lost rapidly upon addition of the PROTAC compared to classical KO systems and at least some downstream consequences occur within hours. Thus, these PROTAC systems allow addressing direct, short-term consequences. It is remarkable that the loss of either RBBP5, ASH2L and DPY30 results in similar consequences, including the fast loss of H3K4 methylation and broad deregulation of gene expression, supporting the view that these proteins are necessary for all KMT2 complexes (22–24).

We have established immortal mouse embryo fibroblasts (iMEFs), in which the endogenous *Ash2l* loci are knocked-out and the cells are rescued by FKBP-ASH2L (24). This fusion protein interacts with dTAG-13, a bifunctional PROTAC, which targets proteins of interest to a cereblon (CRBN)-containing E3 ubiquitin ligase complex (25). The degradation of FKBP-ASH2L occurs within minutes, which is much faster compared to the down-regulation of Ash2l upon KO of the endogenous loci (12,21,24). We hypothesized that the interaction of dTAG-13 through its thalidomide moiety with CRBN might be competed by Lenalidomide, another CRBN ligand (Figure 1A). Indeed, the addition of Lenalidomide results in a rapid re-expression of FKBP-ASH2L, which allows addressing the reversibility of the ASH2L loss consequences.

**Figure 1.**
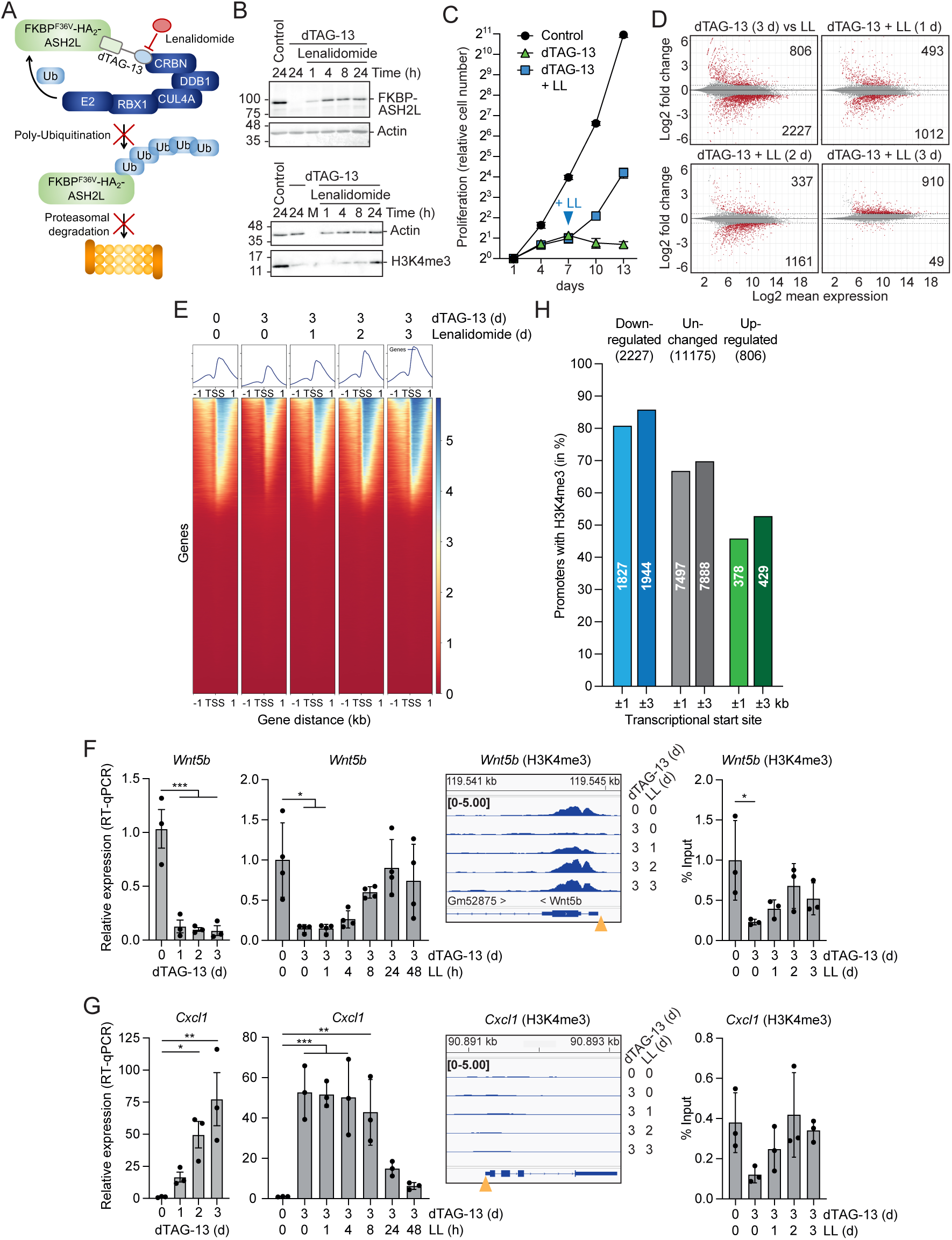
Reversibility of PROTAC induced depletion of ASH2L. A. Schematic summary of dTAG-13 induced degradation of FKBP and antagonize by Lenalidomide. B. iMEF cells expressing the FKBP-ASH2L fusion protein were treated with dTAG-13 (100 nM) and Lenalidomide (50 µM) for the indicated times and lysed in RIPA buffer. ASH2L, H3K4me3 and Actin were analyzed by Western blotting. Uncropped blots are shown in Figure S6. C. Cells were incubated with or without dTAG-13 and Lenalidomide (LL) as indicated and counted every three days (3 technical replicates). D. Cells were treated with Lenalidomide only (LL) or dTAG-13 for 3 days. The latter samples were further incubated for 1, 2 or 3 days with Lenalidomide. Total RNA was extracted and analyzed by 3′ mRNA sequencing. The results are shown as MA plots. Significantly differentially expressed genes are highlighted in red (adj. p-value < 0.05 and logFC > 0.58; two biological replicates, normalized with ERCC spike-in RNA). The total numbers of up- and down-regulated genes are indicated. E. Cells were treated with or without dTAG-13 and Lenalidomide as indicated. ChIP was performed with H3K4me3 selective antibodies. Heatmaps and plot profiles were generated using DeepTools. Normalized signals around transcriptional start sites (TSS) are shown (3 biological replicates). F. Time course of dTAG-13 and Lenalidomide treatment as indicated. RT-qPCR analyses of *Wnt5b* expression (two left panels). IGV Snapshot of the H3K4me3 ChIP-seq data and ChIP-qPCR experiments with a primer pair positioned in the promoter region (yellow arrowhead). Three biological replicates, one-way Anova with multiple comparisons. G. As in panel F, analysis of *Cxcl1*. H. Genes were categorized based on RNA-seq analysis following dTAG-13 treatment for 3 days into upregulated (adj.p.value < 0.05 and logFC > 0.58), downregulated (adjusted p-value < 0.05 and log2FC < −0.58), and unchanged (all remaining genes with baseMean > 10). The promoters of these genes were evaluated for H3K4me3 signals, either in a ±1 kb or in a ±3 kb window relative to the TSS. The proportion of genes showing detectable H3K4me3 signal in each category is displayed in relation to the total number of genes in each group. *p*-values: *≤0.05, **≤0.01, ***≤0.001, ****≤0.0001.

The repression of H3K4me3 and gene expression was largely reversed (21,24,26). Our previous studies indicated that cell cycle progression and proliferation was halted upon depleting ASH2L and that these cells developed a senescence phenotype (21,24,26). Lenalidomide treatment reverted these biological consequences, indicating that cells can recover from extended loss of ASH2L, which includes that senescent cells are reactivated and can enter the cell cycle.

## Results

### PROTAC competition by Lenalidomide reactivates ASH2L expression and cell proliferation

Previously, we established iMEF cells that rely on the expression of FKBP-ASH2L (referred to as ASH2L in the following) for proliferation and evaluated short-term dTAG-13 treatment in these cells (24). Now we extended the analyses to several days and addressed whether the consequences of prolonged ASH2L loss are reversible, both at the molecular and cellular level. Initially, we tested whether the PROTAC-induced knockdown can be reverted by washing out the degradation-inducing compound dTAG-13. Re-expression of ASH2L took several days, indicating that the turnover of dTAG-13 is slow in our cells. Instead, we considered a competition assay, in which the binding of dTAG-13 to CRBN was competed by Lenalidomide. This resulted in a rapid re-expression of ASH2L and, with delay, recovery of global H3K4me3 (Figures 1B and S1A). Moreover, cells kept in dTAG-13 for several days did not proliferate, similar to *Ash2l* KO MEFs (21,24). Lenalidomide addition reactivated cell proliferation within 3 to 4 days, while control cells were not affected (Figures 1C and S1B). These findings suggested that cells can recover from extended loss of ASH2L.

### Reduction of H3K4me3 and altered gene expression are reversed upon re-expressing ASH2L

A major consequence of ASH2L loss is the reduction of H3K4me3 at promoters and the deregulation of many genes by 24 hours (24). After 3 days of dTAG-13 treatment, a time point at which cell proliferation has ceased, H3K4me3 at promoters was repressed and more than 3000 genes were deregulated (Figures 1D, E and S1C). Lenalidomide treatment reversed these effects, but had no effect on control cells. H3K4me3 was reestablished at promoters within 2-3 days, with levels and patterns of this mark being comparable to controls. The number of repressed genes decreased, while a considerable number of genes was upregulated even 3 days after Lenalidomide addition.

This was verified by analyzing individual genes. The repression of many genes occurred within 24 hours while activation developed over 3 days of ASH2L loss (Figures 1F and G and S1). Re-expression of down-regulated RNAs was measurable by 8 to 24 hrs, while up-regulated RNAs were repressed somewhat slower, likely affected by the half-life of the RNAs (Figures 1F and G, and S1). The repressed genes typically lost H3K4me3 at their promoters rapidly, both when measured in ChIP-seq and ChIP-qPCR, an exception being *Flrt1*, which has only background levels of H3K4me3 (Figures 1F and S1D). In contrast, many activated genes had only background levels of H3K4me3, an exception being Rspo2, which has high H3K4me3 with little change upon loss of ASH2L (Figures 1G and S1E and F). For control, H3K4me3 was measured at promoters of genes whose RNA levels were unaffected by loss or regain of ASH2L (Figure S1G). Many showed a strong reduction in H3K4me3, comparable to repressed genes. Finally, we selected two heat shock genes, *Hspa1a* and *Hsph1* (encoding Hsp70 and Hsp105, respectively), which both contain heat shock response elements and are regulated by heat shock factors (27,28). Both genes were efficiently activated to similar levels in cells with and without ASH2L in response to a 42°C heat shock (Figure S1H). The *Hsph1* promoter is marked by broad H3K4me3, while the *Hspa1a* promoter possesses only background levels of H3K4me3 and only small changes were observed upon ASH2L loss (Figure S1I and J). Together, the analysis of individual genes and promoters suggests that the levels of promoter proximal H3K4me3 marks do only partially correlate with expression.

Therefore, we compared the RNA-seq and H3K4me3 ChIP-seq data and addressed the correlation between expression and H3K4me3 levels. Considering a window of ±1 kb relative to the TSS, the majority of down-regulated genes are linked to H3K4me3 positive promoters. This rate was slightly lower for genes that were not deregulated. Of the up-regulated genes less than 50% of the promoters were marked with H3K4me3 (Figure 1H). Slightly more genes were linked to H3K4me3 when a ±3 kb window was analyzed. Thus, down-regulated genes possess preferentially H3K4me3, the majority of unregulated genes are H3K4me3 positive, but it appears that only half of genes that are upregulated are characterized with a promoter proximal region that is H3K4me3 modified.

H3K4me3 at promoters is widely recognized as a hallmark of active promoters (1,29). One of the functions of H3K4me3 at promoters is to recruit and position the TFIID sub-complex of the RNA polymerase II (RNAPII) complex (30–32). Recently, serotonylation of H3Q5 (H3Q5ser) was identified to not only cooperate with H3K4me3, but to be sufficient to recruit TFIID independent of H3K4me3 (33–36). Thus, H3Q5ser may compensate for H3K4me3. We measured H3K4me3Q5ser and H3Q5ser, focusing on promoters that lack H3K4me3 and are linked to up-regulated genes in response to ASH2L depletion. The signals for the H3K4me3Q5ser or H3Q5ser marks were not above background at promoters of up-regulated genes, e.g. *Cxcl1* and *Ccl2* (Figure S1K). At promoters with high H3K4me3, signals for H3K4me3Q5ser were detected, while H3Q5ser was very low. H3K4me3 is linked to open chromatin, which might allow serotonylation of H3Q5 (37). Together, it is unlikely that H3Q5ser compensates for the lack of H3K4me3.

It is possible that gene activation in the absence of H3K4me3 is controlled by transcription factors that do not require this histone mark. Therefore, we analyzed promoter proximal regions for transcription factor binding motifs using Enrichr (38,39). While we identified several transcription factor binding sites (Figure S1L), none of these were statistically significant. Nevertheless, notable are factors known to control cytokine genes, such as C/EBP, NF-κB and PU.1 factors and to regulate proliferation, such as AP1 and SMAD. Thus, it is likely that these genes can be transcriptionally activated by a combination of events, which might include selective histone marks different from H3K4me3 and sequence specific transcription factors.

### Loss of ASH2L results in the induction of senescence

The consequences of *Ash2l* KO are broad, which include the deregulation of genes, a proliferation stop, and the induction of a senescence-like phenotype (21). Because senescence is a heterogenous process dependent on the cell type, the molecular trigger and the wiring of downstream pathways (40–44), we wanted to obtain additional evidence for senescence in response to ASH2L loss. One criterium is a proliferation arrest, as also seen in our iMEF cells upon depleting ASH2L (Figure 1C), which is thought to be permanent (41,45). Thus, the reactivation of cell proliferation shown above suggests that at least some cells are not permanently arrested, possibly because they are not senescent. One marker of senescence is the increase of senescence-associated β-galactosidase (SA-β-Gal) activity (42,46). After 3 days of dTAG-13 treatment, the majority of cells stained positively for SA-β-gal (Figure 2A and B). As expected from reestablishing proliferation upon re-expression of ASH2L, the number of SA-β-gal positive cells decreased over time (Figure 2B).

**Figure 2.**
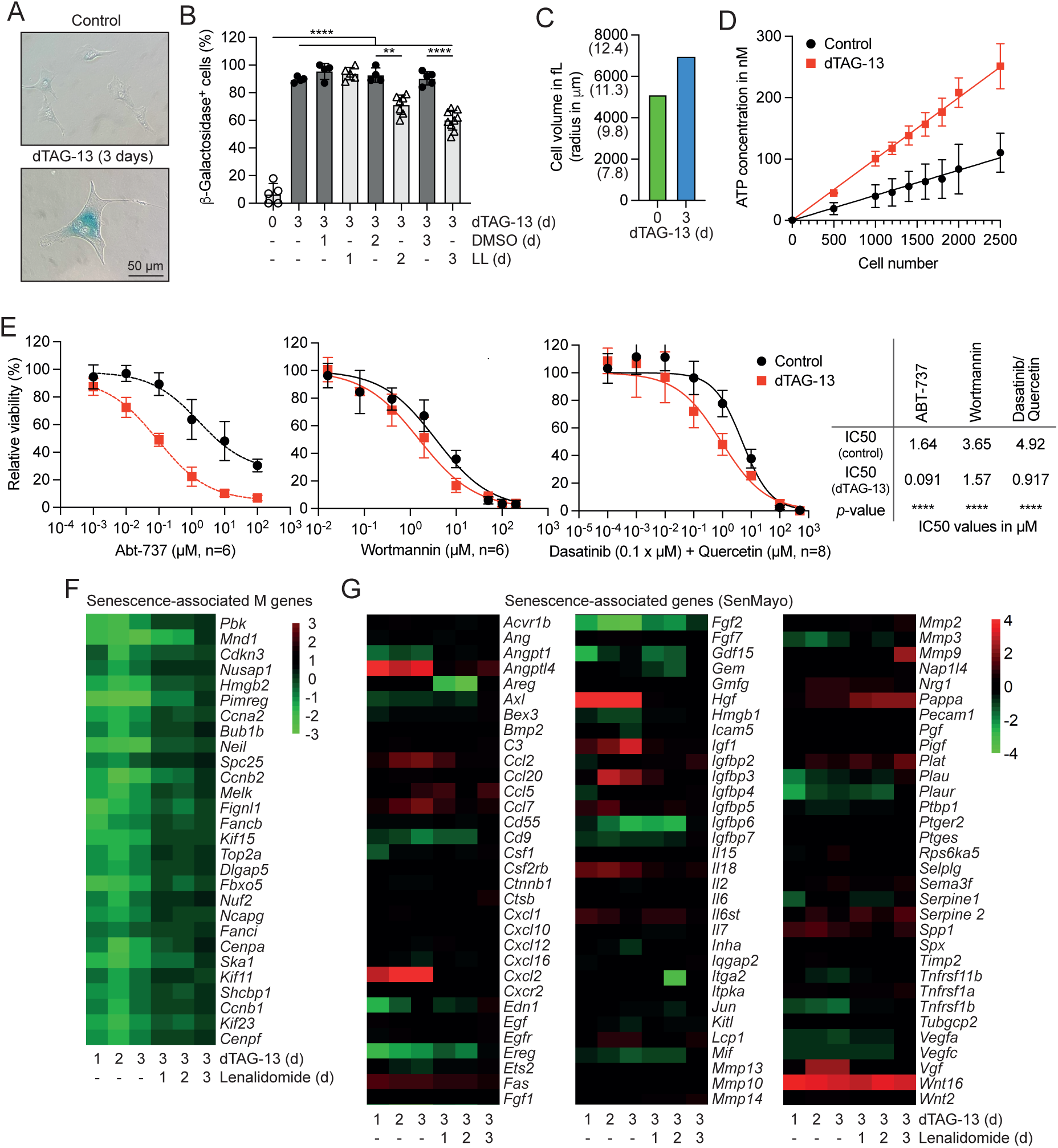
Loss of ASH2L induces senescence. A. iMEF cells expressing the FKBP-ASH2L fusion protein were incubated with ±dTAG-13 for 3 days. Cells were fixed and stained for SA-β-galactosidase activity (blue). B. iMEF cells were incubated with or without dTAG-13 and Lenalidomide (LL) for the indicated times. The cells were fixed and stained for SA-β-galactosidase activity. Percentages of β-gal positive cells were determined. Mean values ± SD of 5-9 measurements. Statistical analysis with one-way Anova/multiple comparisons. C. Cells were treated ±dTAG-13 for 3 days. Average cell size was determined on a Casy cell counter (peak size volume is indicated). D. Cells were treated ±dTAG-13 for 3 days, then seeded at indicated cell numbers in 96-well plates. ATP concentration was determined using the CellTiter Glo Assay compared to an ATP standard curve. Shown are mean values ± SD of 3 experiments. E. Cells were treated ±dTAG-13 for 3 days. Subsequently senolytic compounds at the concentrations indicated were added for 24 hrs, washed and cultured for an additional 48 hrs. Relative viability was determined using the CellTiter Glo Assay. Shown are mean values ± SD of 6 to 8 experiments. Sigmoidal fitting curves were added using the GraphPad Prism Software and constrained at 0% (bottom) and 100% (top). Resulting IC50 values for each curve and p-values using nonlinear fit are summarized in the table. F. The expression of M genes (Ref (21)) were analyzed from RNA-seq data in response to dTAG-13 and Lenalidomide treatment as indicated. G. As in panel F, analyzing the SenMayo genes (Ref (63)). *p*-values: *≤0.05, **≤0.01, ***≤0.001, ****≤0.0001.

Senescence is associated with increased cell size (46–48). Morphologically the dTAG-13 treated iMEF cells appeared larger and flat (Figure 2A). The cell size increased by 25% upon dTAG-13 treatment for 3 days (Figure 2C). Additionally, we measured the ATP content, expecting higher levels in senescent cells due to increased cell size, although senescent cells have reduced ATP levels (49,50). ATP levels roughly doubled upon loss of ASH2L (Figure 2D), consistent with increased cell size, as also further argued below. Moreover, senescent cells have been described to be particularly sensitive to senolytics that target the mitochondrial apoptosis pathway (51). We employed Abt-737, a BH3 mimetic, which inhibits the anti-apoptotic Bcl-2 and Bcl-xL, two members of the Bcl-2 family(52–55). Cells with depleted ASH2L were roughly 20 times more sensitive to Abt-737 compared to control cells (Figure 2E). Senescent cells were also more sensitive to Wortmannin, a senolytic that inhibits the PI3 kinase pathway (Figure 2E) (56). Finally, the combination of Dasatinib and Quercetin is preferentially targeting senescent cells (21,57,58). Again, dTAG-13 treated cells were more sensitive (Figure 2E). Together, these observations, i.e. inhibition of cell proliferation, SA-β-gal positivity, increase in cell size, and sensitivity to senolytics, support the interpretation that the loss of ASH2L promotes senescence.

### Reactivation of ASH2L reverts senescence markers

One of the hallmarks of senescence is the activation of SASP (senescence-associated secretory phenotype) genes, a heterogenous set of genes being controlled in distinct settings. The secreted factors promote inflammation and affect aging and tumor formation (41,44,59,60). We have argued previously that this may be difficult to observe as the loss of ASH2L and subsequently of H3K4me3 at promoters may prevent gene activation (21). Therefore, we defined a set of down-regulated genes as marker of senescence, referred to as M genes, which were also repressed in other senescent cells (21,61,62). These were repressed upon ASH2L depletion and were reactivated upon addition of Lenalidomide (Figure 2F). In addition, we evaluated a set of genes recently reported to be senescence-associated, referred to as SenMayo (63). We noted that the response of these genes was heterogenous in our cells (Figure 2G). Some SASP genes listed in SenMayo, including *Cxcl1*, *Ccl2*, and *Angptl4*, are activated upon loss of ASH2L and subsequently repressed by Lenalidomide (Figures 1G and S1E). Together, these findings suggest that senescence-associated altered gene expression, either activated or repressed upon dTAG-13 treatment, is reversed upon Lenalidomide treatment.

Despite these observations we did not observe terms relating to senescence in the GO assessment of our RNA-seq data (Figure S2). Nevertheless, associated with down-regulated genes after loss of ASH2L were terms relating to cell proliferation, cell division, and replication. In particular the terms relating to G2/M transition and mitosis are consistent with our observations in the hematopoietic system upon KO of *Ash2l* (Luscher-Firzlaff), and with the rapid proliferation arrest upon ASH2L depletion (Figure 1). In contrast, when up-regulated genes were evaluated, terms associated with cell migration and with extracellular matrix organization were evident, processes that are typically activated in senescent cells (59). These patterns changed rapidly upon reactivation of ASH2L. Down-regulated genes were now linked to migration, angiogenesis, and wound healing, while up-regulated genes were associated again with migration but also with cell differentiation (Figures S2B-D). We also analyzed the GO terms for biological processes of up-regulated genes, whose promoters were either positive or negative for H3K4me3 (Figure S2E and F). Again, many processes were linked to migration, differentiation, and extracellular matrix organization, which are all deregulated in senescent cells. However, terms like senescence, DNA damage or stress response were not found in the top group, although they appeared with low adjusted p-values.

### Senescent cells can re-enter the cell cycle

The findings so far suggest strongly that the loss of ASH2L induces senescence. This is reversed by re-expressing ASH2L, which results in cell cycle entry and proliferation. However, a small percentage of cells did not show enhanced staining for SA-β-gal, which we used as primary determinant of senescence (Figure 2B). Thus, we cannot exclude that the recovery of cell proliferation is due to those few cells that are SA-β-gal negative. Therefore, we generated cells that express histone H2B-mCherry to be able to follow individual cells microscopically. Treatment with dTAG-13 was conducted for 4 days and the cells stained for SA-β-gal using a life cell stain (C12FDG). The cells with high SA-β-gal signals were then sorted according to cell size into four subpopulations P1-P4, which represent low, medium, high and very high cell size, respectively (Figure S3A). An aliquot of each population was fixed and stained for SA-β-gal. In all four sorted subpopulations the number of SA-β-gal positive cells were larger than in the unsorted population (Figure 3A). On average, the cells in the very high (P4) and high (P3) populations were larger than the unsorted, middle and low populations, and all sorted populations were larger than untreated cells (Figure 3B). The unsorted cells roughly doubled their size upon loss of ASH2L, and thus further increased their size compared to 3 days of treatment (Figure 2C). Of note, the very high and high populations consisted of significantly more senescent cells with over 97% and 96% SA-β-gal positive cells, respectively (Figure 3A), and, furthermore, the very high population was almost 3 times as large as control cells, suggesting that these populations were highly enriched for senescence cells.

**Figure 3.**
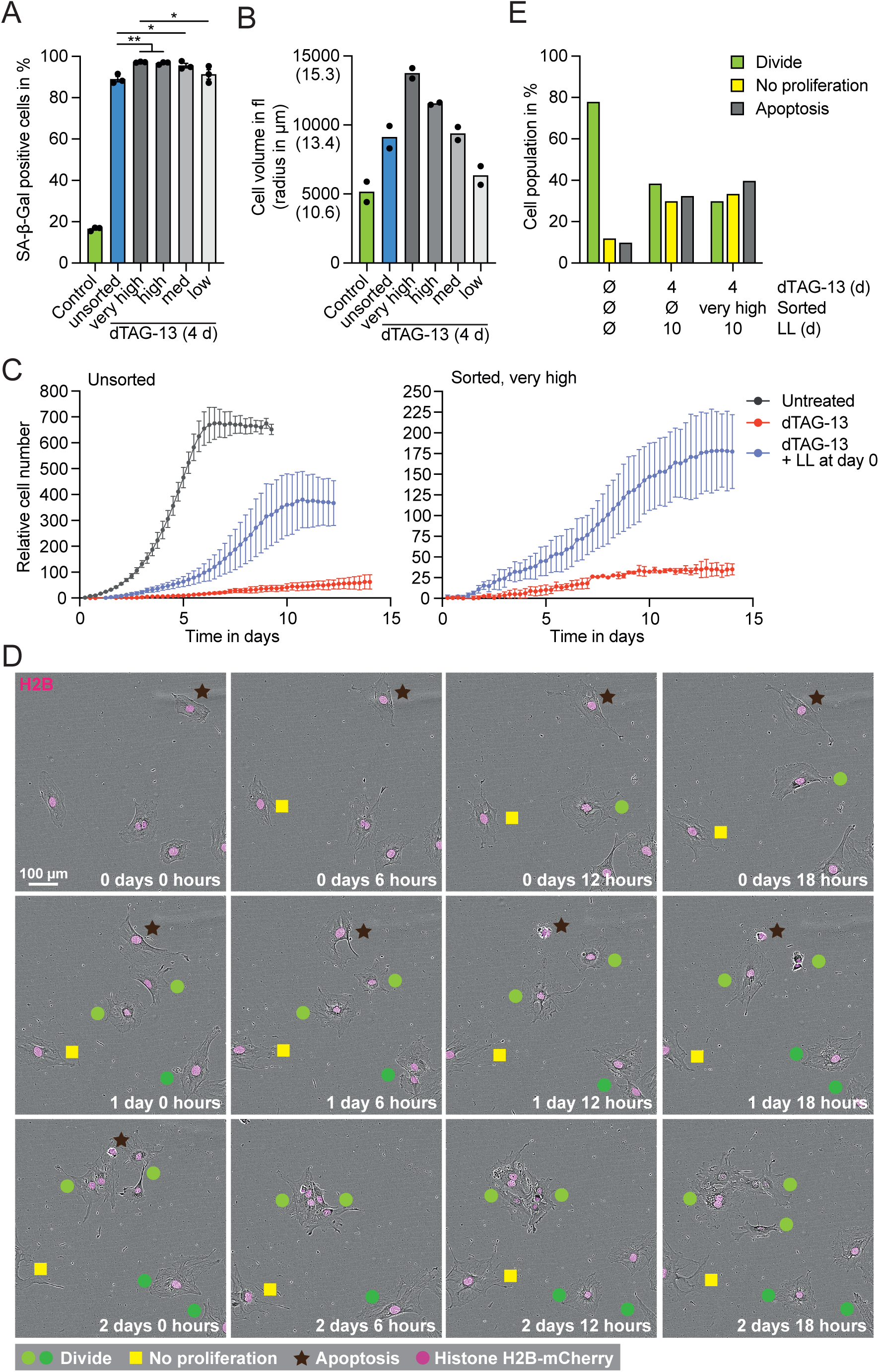
Re-expression of ASH2L reactivates senescent cells. A. iMEF cells were treated with or without dTAG-13 for 4 days. The cells were FACS sorted according to SA-β-gal positivity (using the life cell stain C12FDG) and cell size (forward-scatter). Samples of each group as well as unsorted control and dTAG-13 treated cells were fixed and stained for SA-β-gal activity using X-gal. Percentage of positive cells was determined in one experiment by three independent people (data points). One-way Anova with multiple comparisons. B. Cells were treated and FACS-sorted as described for panel A. Cell size of each population was determined in two independent experiments on a CASY Cell Counter. C. dTAG-13 treated cells were FACS sorted, replated ±Lenalidomide, and proliferation was documented in the Incucyte SX5 Live-Cell analysis system over 14 days at time intervals of 6 hours. Shown are the growth curves of unsorted control cells and sorted cells of the very high group (see Figure S3A). Shown are mean values ± SD of three replicates. D. Microscopic pictures of the time course of cells of the high group. Indicated are cells that divide, enter apoptosis, or remain senescent in the first 66 hrs. The cells express histone H3B-mCherry (in pink). E. Cells obtained in the Incucyte over 10 days were visually analyzed for their ability to proliferate or not, or to undergo apoptosis. p-values: *≤0.05, **≤0.01, ***≤0.001, ****≤0.0001.

The sorted and unsorted cells were then followed microscopically at the single cell level in the continuous presence of dTAG-13 or upon treatment with Lenalidomide. We noticed that ASH2L depleted cells started to proliferate in the presence of Lenalidomide, while dTAG-13 only cells did not (Figure 3C). The four sorted populations differed as the cells in the low group started to proliferate in the presence of dTAG-13 without Lenalidomide treatment, as did unsorted cells, while the cells in the very high, high and medium groups did not (Figures 3C, S3B and C). In response to lenalidomide, the cell numbers reached were lowest in the very high population, likely because these cells were the largest at the beginning of the analysis and, being contact inhibited, reaching confluency earlier. Note that because of the differences in cell size of the starting populations, the analysis of cell numbers and confluency results in distinct curves (Figures 3C, S3B and C).

To analyze cell recovery in more detail, several hundred cells were followed over 10 days (Figures 3D and S3E). We identified cells that either proliferated, died, or remained large and flat. Quantification revealed that about a third of the cells remained large and did not proliferate, both in the total as well as in the very high sorted population of cells (Figure 3E). In addition, about a third of the cells died, while another third divided at least once. Some of the cells divided more than once, about 18% and 10% in the unsorted and very high populations, respectively (Figures 3D and S3D). In control cells, not treated with dTAG-13, the number of either non-proliferating or apoptotic cells was in the range of 10%. Thus, despite similarities in cell size and morphology, SA-β-gal staining, and lack of proliferation, three distinct subpopulations are apparent once ASH2L is reactivated. That cells can divide, die or remain senescent suggests that the loss of ASH2L in our iMEF cells generates heterogeneity as has been reported in cell populations previously (40). Importantly, at least some of the senescent cells can be reactivated to enter the cell cycle and to proliferate.

### Molecular characterization of reactivated senescent cells

The findings above indicate that, upon loss of ASH2L, the large majority of cells have characteristics of senescent cells, defined by SA-β-gal positivity, cell cycle arrest, cell size, and an altered gene expression program. These cells are heterogeneous as some can re-enter the cell cycle and contribute to the proliferative response upon addition of Lenalidomide, but obviously not all large, SA-β-gal positive cells divide again (Figure 3D and E, and SE3). Therefore, we performed single cell RNA-seq experiments to determine whether the observed functional heterogeneity can be visualized by distinct gene expression programs.

We generated cells treated with and without dTAG-13 and/or Lenalidomide (diagramed in Figure 4A). The RNAs of individual cells of the different samples were sequenced using Parse Biosciences Evercode WT platform. Principal component (PC) analysis of the obtained data sets showed that control cells and Lenalidomide only treated cells, either for 24 or 96 hrs, colocalized (Figure 4B, groups 1-3), supporting the findings above that Lenalidomide did not influence cell proliferation and gene expression (Figures 1 and S1). dTAG-13 treatment provoked major shifts by 24 and 48 hrs with no further change by 72 hrs (Figure 4B, groups 4-6). Interestingly, already after 4 hrs, and more pronounced after 24 hrs of Lenalidomide treatment, the gene expression program changed compared to 48 and 72 hrs dTAG-13 samples. Lenalidomide treatment longer than 24 hrs did not result in additional major changes. Of note is that the Lenalidomide rescued cells (Figure 4B, groups 8-11) did not colocalize with control cells (Fig. 4B, groups 1-3). This might reflect the above-described functional heterogeneity.

**Figure 4.**
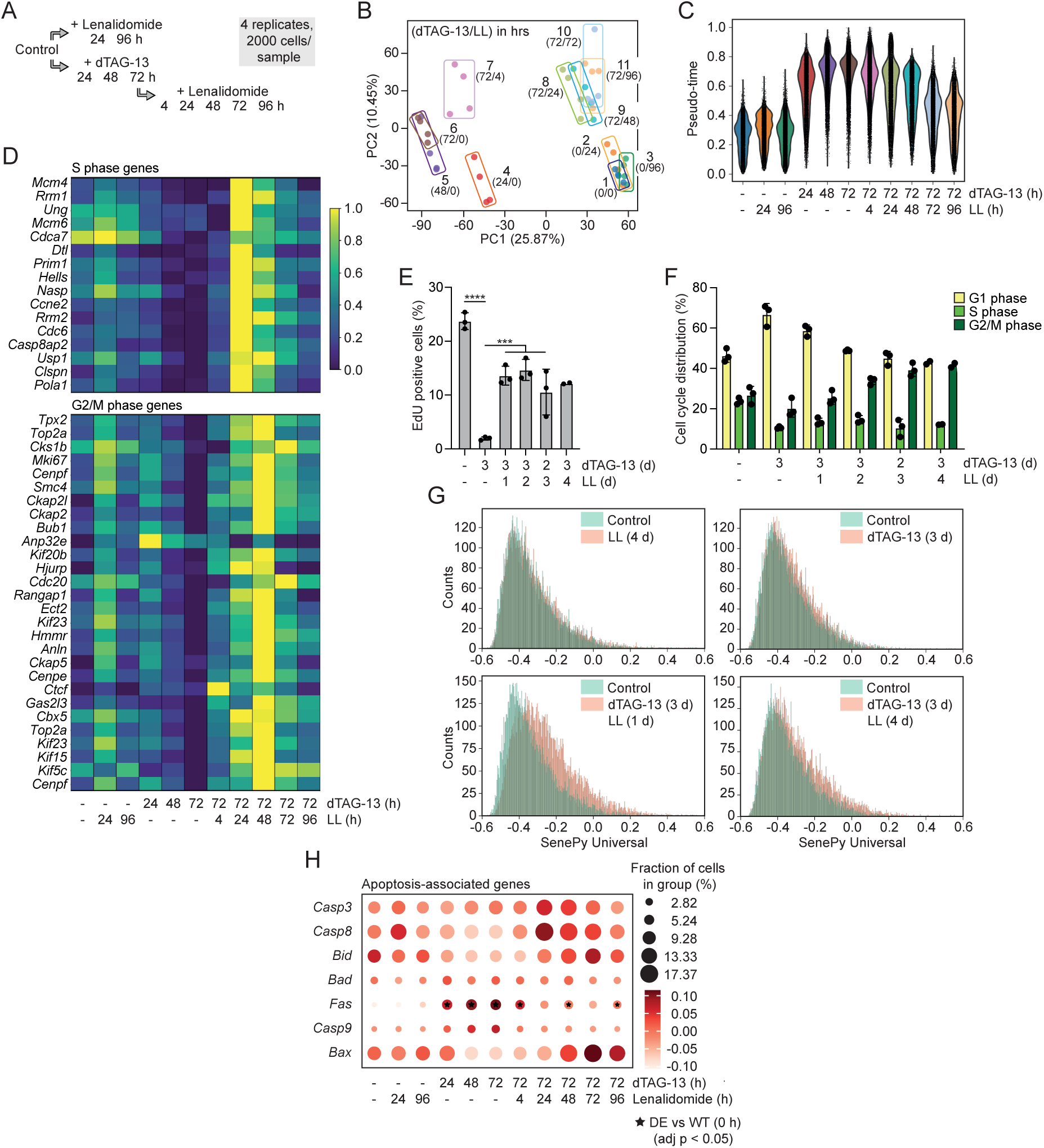
Re-expression of ASH2L induces a transient SASP phenotype. A. Schematic summary of the samples generated for the scRNA-seq analysis. B. Principal component analysis of the scRNA-seq data set. Indicated are the 11 samples (in quadruplicates) treated ±dTAG-13 in hours/±Lenalidomide in hours. The quadruple samples are boxed. C. Pseudo-time analysis of the samples treated with and without dTAG-13 and Lenalidomide as indicated. D. Heatmap of genes associated with S phase and with G2/M phase of the cell cycle in response to dTAG-13 and Lenalidomide treatment. E. Cells were treated with dTAG-13 and Lenalidomide as indicated. The cells were labeled with EdU for 3 hrs priot to harvesting. The cells were fixed, stained for EdU, incubated with propidium iodide (PI) and analyzed by flow cytometry. Three biological replicates, one-way Anova/multiple comparisons. F. The PI signals of the experiments used for panel E were analyzed separately. The distribution of G1, S, and G2/M phase DNA contents are summarized. G. The senescence-associated genes in SenePy Universal (Ref (66)) were evaluated in the scRNA-seq data set of the indicated time points. H. The expression of genes associated with apoptosis were analyzed in the scRNA-seq data set. p-values: *≤0.05, **≤0.01, ***≤0.001, ****≤0.0001.

We also applied high-resolution pseudo-time ordering of cells, originally designed to assess the differentiation of hematopoietic cells (64). This analysis supported our previous analyses. Lenalidomide treatment had little effect, while 24 hrs of dTAG-13 addition provoked a major shift in the pseudo-time (Figure 4C). This was further accentuated at later time points and reverted upon Lenalidomide competition. The cells remained more heterogeneous than the control population, again in line with the three subclasses of cells observed during Lenalidomide rescue (Figure 3D and E). However, despite the differences observed between control cells and Lenalidomide rescued cells, both in the PC and the pseudo-time analyses, we were unable to distinguish subgroups of cells that would explain the three functional cell populations.

Because cell cycle arrest was a prominent and rapid effect upon loss of ASH2L (Figure 1C), we analyzed the expression of genes associated with functions in S phase and G2/M phase using the scRNA-seq data. These genes were downregulated by 48 and 72 hrs of dTAG-13 treatment and reactivated quickly in response to Lenalidomide (Figure 4D). By 4 hrs the expression of many of these genes increased, by 24 hrs S phase genes and by 48 hrs G2/M phase genes were transiently induced (Figure 4D). This is consistent with reassuming proliferation and increasing cell numbers by 2 to 3 days of Lenalidomide addition (Figure 1C), and with the transient increase in the overall S and G2/M phase scores (Figure S4A). Also supporting the transient cell cycle arrest is that many Myc target genes were down-regulated in ASH2L depleted cells and reactivated upon Lenalidomide treatment (Figure S4B). Moreover, the rapid re-entry into the cell cycle is documented by the incorporation of the nucleotide analog 5-ethynyl-2’-deoxyuridine (EdU). In dTAG-13 treated cells, the incorporation of EdU was halted but recovered upon Lenalidomide treatment (Figures 4E and S4C). Of note, the propidium iodide (PI) stained dTAG-13 treated cells accumulated only partially in G0/G1, with a substantial number of cells possessing S and G2/M phase DNA contents (Figures 4F and S4C). This is consistent with our previous observation of a cell cycle block upon KO of *Ash2l* (21). This also argues against an accumulation of the cells in a quiescent state, which is fundamentally different from senescence (41,65). The number of S phase cells did not recover to the level of control cells, while G2/M phase cells increased upon Lenalidomide treatment (Figure 4F), in agreement with a subpopulation of cells reentering the cell cycle.

Next, we addressed the expression of genes that are associated with senescence. Our data from the bulk RNA-seq and the qPCR experiments were heterogeneous. For example, only few genes of the SenMayo gene set were regulated in response to ASH2L modulation (Figure 2G) (63). Therefore, we analyzed the SenePy Universal gene set, which has been linked recently to senescence in multiple cell types (66). We observed little change in gene expression after three days of dTAG-13 treatment (Figure 4G). However, these genes were activated in response to 24 hrs Lenalidomide treatment, an effect that was transient. A similar observation, albeit less pronounced, was made when the SenMayo gene set was analyzed (Figure S4D). Many chemokines, interleukins, growth factors and proteases are induced during senescence as part of the SASP response (67). Many of these are deregulated during dTAG-13 treatment and a subset is activated once ASH2L expression is reestablished, similarly to the SenePy and SenMayo data sets (Figure S4E). It is possible that many of these genes cannot be induced in the absence of ASH2L and when promoter-associated H3K4me3 is lacking, although the relevant signals are present. Once ASH2L is re-expressed, the signals that promote the expression of these senescence-associated genes are still active and, together with reestablishing the positive H3K4me3 mark, are now promoting their expression. Once the senescence signals vanish, the expression of these genes approaches control values.

A considerable number of cells die upon re-expression of ASH2L (Figure 3D and E). The analysis of genes associated with apoptosis demonstrates that many are induced in the recovering cells (Figure 4H). In particular, *Bid* and *Bax* are induced in a subpopulation of Lenalidomide treated cells. These two genes, which encode proapoptotic proteins (68), may explain the increased apoptosis observed in some of the cells, in which dTAG-13 is competed with lenalidomide. Thus, although the scRNA-seq data did not reveal the expected three distinct subpopulations, the gene expression patterns support these phenotypes. It suggests that the gene expression patterns are overlapping and the cellular outcome, i.e. proliferation, cell death, and remaining senescent, is determined by potentially small differences or fluctuations in gene expression during recovery.

### Senescence induced by loss of ASH2L is the consequence of KMT2 complex failure

It is unclear whether all the effects observed in response to ASH2L loss are due to its function in KMT2 complexes. Of note is that the WRAD component WDR5, which, similar to ASH2L, has a key role in KMT2 complex functions, interacts with various transcription factors, and can read H3K4me2 and H3Q5ser (12,69). Furthermore, WDR5 is part of for example histone acetyltransferase complexes and associates with the anaphase promoting complex, activities that go beyond the functions attributed to KMT2 complexes (69,70). Considering these multiple functions of WDR5, we addressed whether additional activities can be predicted for ASH2L. We took advantage of its well-defined, conserved domains (Figure 5A). The SPRY and SDI domains function to recruit RBBP5 and DPY30, respectively, two partner proteins in the KMT2 complexes (71–76). The HWH domain can interact with DNA (77,78), while the role of the unstructured N-terminal and of the atypical PHD domains are not fully understood. Therefore, we generated ASH2L mutants that lack individual domains (Figure 5A). These mutants were stably introduced as GFP-fusion constructs into the FKBP-ASH2L cells. All constructs showed comparable levels of expression of the different proteins (Figure 5A). To address which domain(s) are important for cell proliferation, ASH2L was depleted with dTAG-13 and the GFP-fusion proteins induced using doxycycline. In control cells, transduced with an empty lentivirus, the cells did not proliferate in the absence of ASH2L (Figure 5B and C). Both the SPRY and the SDI domains are important for proliferation. In addition, a small effect was observed upon deletion of the PhD domain (Fig. 5B and C). These findings suggest that cell proliferation is mainly dominated by the ability to interact with RBBP5 and DPY30 and thus to form KMT2 complexes.

**Figure 5.**
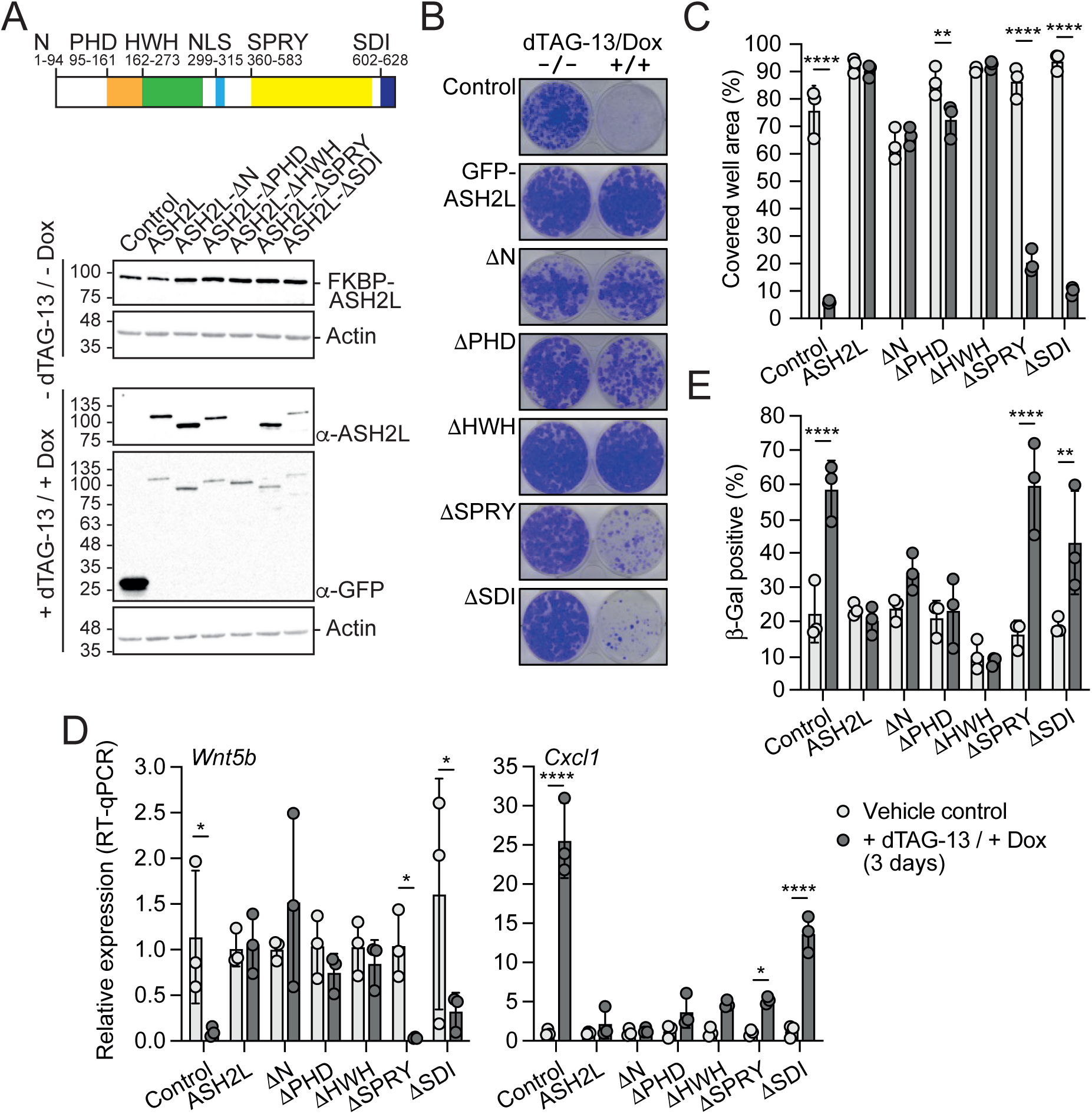
ASH2L domains necessary for KMT2 complex formation are critical for proliferation and senescence. A. Scheme of full-length ASH2L. Major domains are indicated, which are deleted individually in the different ASH2L mutants. Their expression is analyzed on Western blots. Please note that the ASH2L antibody used recognizes epitopes in the HWH domain. B. Cells were treated with or without dTAG-13 and Doxycycline to induce loss of FKBP-ASH2L and expression of ASH2L and ASH2L mutants, respectively. Cell proliferation was assessed by plating 2000 cells/6 cm plate and staining the cells 6-8 days later using Crystal Violet. C. Summary of cell proliferation assays as in panel B. Three biological replicates, one-way Anova/multiple comparisons, mean values ± SD. D. Cells were treated with or without dTAG-13 and Doxycycline for 3 days. The RNA of the indicated genes was analyzed using RT-qPCR. Three biological replicates, one-way Anova/multiple comparisons, mean values ± SD. E. Cells were treated with or without dTAG-13 and Doxycycline for 3 days and stained for SA-β-gal activity. Three biological replicates, every replicate was counted blinded by 3 persons, one data point is the mean value of the three countings. Shown are mean values ± SD, one-way Anova/multiple comparisons. p-values: *≤0.05, **≤0.01, ***≤0.001, ****≤0.0001.

The different deletion mutants were further analyzed regarding gene expression. We evaluated three repressed (*Wnt5b*, *Lin7a* and *Mcm6*) and two activated (*Cxcl1* and *Ccl2*) genes upon ASH2L loss. In these newly established stably transduced cells, *Wnt5b* and *Lin7a* were efficiently repressed in response to dTAG-13 treatment, while the effect on *Mcm6* was less pronounced (Figures 5D and S5). As for cell proliferation, the SPRY domain and the SDI domain were relevant for rescuing gene expression. The other domains appeared dispensable as deletions of either the N-terminal region, the PHD domain or the HWH domain did not rescue gene expression. The two activated genes, *Cxcl1* and *Ccl2*, were partially induced when the SPRY domain or the SDI domain mutants were expressed (Figures 5D and S5). Thus, ASH2L mutants lacking either the SPRY or the SDI domain cannot efficiently repress the dTAG-13 induced loss of ASH2L. Because *Cxcl1* and *Ccl2* are SASP genes, we expected that the SPRY and SDI domains are also relevant for the induction of senescence. Indeed, dTAG-13 treated cells that express ASH2LΔSPRY or ASH2LΔSDI, but not any of the other mutants, showed increased SA-β-gal activity. These findings together with the effects on proliferation suggest that both the SPRY and the SDI domain are critical for the function of ASH2L, arguing that the observed phenotypes upon depletion of ASH2L are due to the formation of KMT2 complexes and thus likely to altered H3K4 methylation.

## Discussion

ASH2L is a core subunit of KMT2 COMPASS-like complexes, which associate with a large number of transcription factors to mediate H3K4 methylation (12). Depleting ASH2L results in loss of H3K4me3 at and reduced accessibility of promoters, and broad deregulation of gene transcription, after both KO of endogenous loci and depletion of ASH2L using a PROTAC system (21,24,26). Here we expanded our analysis of the PROTAC system by demonstrating that the effects of ASH2L loss are largely reversible by competing the binding of dTAG-13 to a CRBN-containing E3 ubiquitin ligase complex using Lenalidomide. This results in the rapid re-expression of ASH2L, the reappearance of H3K4me3 at promoters, and gene expression is largely reversed. Moreover, part of the cells re-enter the cell cycle, proliferate and overcome senescence. This correlates with the recovery of gene expression linked to these biological processes. Moreover, the analysis of ASH2L deletion mutants suggests that the molecular and biological consequences are mainly due to the function of ASH2L in KMT2 complexes.

The analyses of RNA expression, in both bulk and the single cell studies, demonstrate large changes upon depletion of ASH2L. This occurs rapidely and more than 3000 RNAs are deregulated 3 days after dTAG-13 addition (Figures 1 and 4). Many of these changes are reversed rapidly. We note that in the principal component and pseudo-time analysis of the scRNA-seq data set a change is already visible 4 hrs after the addition of Lenalidomide (Figure 4). At this time point, ASH2L is re-expressed but H3K4me3 has not recovered yet (Figure 1). At later time points, gene expression approaches control levels, but clear differences are observed. In the bulk RNA data, many genes are up-regulated, while the number of down-regulated genes is very low after 3 days of Lenalidomide treatment. Similarly, the scRNA data show a clear difference to control cells or cells treated with Lenalidomide only (Figure 4). One interpretation is that the rescued cell population is heterogenous, as discussed below, which results in distinct RNA expression patterns that support the functional differences.

Cellular senescence is a response to many different forms of stress, including telomere attrition, the activation of oncogenes, and chemotherapy (41,79–81). Resuming proliferation of a senescent cell population upon re-expression of ASH2L was unexpected. While reentry into the cell cycle can be documented at the population and single cell level, by the activation of S and G2/M phase genes, and the incorporation of EdU upon Lenalidomide treatment, senescence is more difficult to define. This is because senescence is a heterogeneous phenotype and is typically referred to as irreversible (41,46). Therefore, it was important to analyze multiple markers, in addition to SA-β-gal positivity and cell cycle arrest. Increased cell size has been linked to senescence (42,48). Excessive cell growth results in altered ratios of cellular proteins that interfere with cell proliferation and supports senescence (82,83). Preventing cell cycle progression in the presence of growth factors and an increase in cell size can induce senescence (84–87). We observed that on average the cell size doubled after 4 days of dTAG-13 treatment, with the top group of SA-β-gal positive cells nearly tripling their size (Figures 2 and 3). Moreover, compared to control cells, senescent cells are more sensitive to senolytics, which are assessed in clinical trials to treat multiple diseases and age-related conditions in humans (42,88–91). Indeed, our dTAG-13 treated cells showed increased sensitivity to different senolytics, further supporting the interpretation that ASH2L loss provokes senescence (Figure 2E).

An important characteristic of senescence is the activation of the SASP gene expression program that includes chemokines, cytokines, growth factors and proteases. Typically, this drives a pro-inflammatory environment that contributes to processes such as aging but can also promote cancer progression (41,46,92). SASP is highly heterogeneous, depending on the stimuli applied and the cell type studied (42,43,93). We did not observe a classical SASP phenotype in our cells. However, our gene expression studies addressing the down-regulated M genes (21,61,62), the SenMayo and the SenePy datasets (63,66), and the analysis of selected SASP genes such as *Cxcl1* and *Ccl2* provide a strong link to senescence. We have argued before that many SASP genes may not be activated in the absence of H3K4me3 at promoters (21), a possibility being that the reduction of H3K4me3 at promoter reduces their accessibility, which may affect transcription factor and RNAPII complex binding (26). However, promoters that lack H3K4me3 may not be sensitive to ASH2L loss. Therefore, a possible interpretation is that the loss of ASH2L generates stress signals that can only partially be translated into gene expression. But once ASH2L expression is reestablished, additional senescence-associated genes are activated transiently, as documented for the SenePy and SenMayo data sets (Figures 4G and S4D). Because also cell cycle-associated genes are activated, this may result in a conflict of signals leading to apoptosis in some of the cells. Indeed, in a subpopulation of cells several pro-apoptotic genes are induced upon re-expression of ASH2L (Figure 4H).

Together, the combination of the different functional and molecular changes upon loss of ASH2L in our iMEF cells, which include cell cycle block, proliferation arrest, SA-β-gal positivity, increase in cell size, sensitivity to senolytics, and changes in gene expression programs, are consistent with efficient induction of senescence. Importantly, this is to a large degree nullified when the PROTAC dTAG-13 is competed with Lenalidomide.

The cellular response is heterogenous during the recovery upon reactivating the expression of ASH2L. How does reentry into the cell cycle and proliferation relate to the common assumption that senescence is irreversible? It is of note that several reports have suggested that under specific conditions senescence can be reverted. Early on it was demonstrated that proliferation can be stimulated in human replicative senescent fibroblasts upon expression of the multifunctional SV-40 large T antigen, a viral protein that can inactivate both the p53 and the RB pathways (94), while certain cellular oncoproteins promote senescence to prevent cell transformation (59). Moreover, the expression of Oct4, Sox2, Klf4 and Myc, the so-called Yamanaka factors that can reprogram differentiated cells (95), ameliorate age-associated characteristics, including the repression of senescent marker genes and SA-β-gal activity (96). Also, a complex of the scaffold attachment factor SAFA and the lncRNA PANDA promotes senescence when interacting with polycomb repressive complexes, which can be reverted by depleting PANDA (97). Another example relates to chemotherapy-induced senescence in a B cell lymphoma model. Manipulating either the heterochromatin marker H3K9me3 or the tumor suppressor p53 can reactivate cells to enter the cell cycle. Interestingly, these cells possess stem cell properties with a higher tumor initiating potential compared to the starting population (98). Relating to these findings, it has been suggested that senescence is a potential mechanism to allow long-term survival of dormant tumor cells that can result in relapse upon their reactivation (99,100) (101,102). Thus, understanding how senescent cells can be reactivated to proliferate is of broad interest.

The findings discussed above also indicate that changes in gene expression and chromatin reorganization, possibly due to differential expression of transcription factors and epigenetic regulators, have the potential to induce or even reverse senescence (103,104). For example, an increase in cryptic transcription has been noted as a feature of senescent cells (105,106). This results from transcriptional initiation in gene bodies and potentially generates non-functional or even toxic products. Mechanistically, this is associated with a decrease of H3K36me3 in gene bodies, a histone mark which prevents H3K4me3 and enhances local DNA methylation by recruiting the H3K4 demethylase KDM5B and the DNMT3B DNA methyltransferase, respectively (107,108). While cryptic transcription may not be efficient due to the loss of KMT2 complexes in our system, transcription factors such as AP1 family members and PU.1 have been suggested to be relevant (105,109). Binding sites for these factors are linked to upregulated genes upon loss of ASH2L that do not possess H3K4me3 at their promoters (Figure S1L). Interfering with AP1 factor functions can revert senescence (109).

Together, our findings provide evidence that ASH2L loss induces cellular stress that results in cellular senescence. The analysis of ASH2L mutants suggests that this is the consequence of KMT2 complex inhibition and thus reduction or loss of H3K4 methylation marks. This represses many genes, including some that encode SASP factors. Other SASP genes are activated, likely because they do not depend on H3K4me3 at their promoters for being transcribed. This indicates that deregulating KMT2 complexes can affect a cell’s decision whether to proliferate or to enter a senescent state, and to reenter the cell cycle, which is potentially relevant for controlling age-associated phenotypes, including the reactivation of dormant tumor cells.

## Material and Methods

### Tissue culture

Immortalized mouse embryo fibroblast, in which the endogenous *Ash2l* loci were deleted and which express from a stable transgene FKBP-HA2-ASH2L from a stably introduced transgene (iMEF clone NG3), have been described previously (24). iMEF cells and lines derived from these cells and HEK293T cells were cultured in DMEM supplemented with 10% (v/v) fetal bovine serum (FCS) and 1% Penicillin/Streptomycin (v/v) at 37°C in a humidified incubator at 5% CO2.

iMEF cells expressing mCherry-H2B or different ASH2L mutants were generated through lentiviral infection. For virus production, a third-generation packaging system was transfected into HEK293T cells, including three helper plasmids (pMDLg/pRREm pRSV-rev and pLP-VSV-G), with a pLentiLox-mCherry-H2B or pLEX-305-N-EYFP plasmids expressing ASH2L mutants. Culture supernatants from the transfected HEK293T cells were collected after 48 hrs and passed through a 0.45 µm PVDF filter. iMEF cells were seeded at a confluency of roughly 40% and incubated with lentivirus-containing supernatant diluted 1:2 with fresh medium and supplemented with polybrene (8 µg/mL) for 6 hours. Transduction was repeated 24 hours later. High mCherry-H2B expressing cells were sorted by FACS. Cells expressing ASH2L mutants were selected in Hygromycin (20 µM/mL).

To deplete FKBP-HA2-ASH2L, cells were treated with dTAG-13 (100 nM). Competition experiments were carried out by adding Lenalidomide (50 µM) to the culture medium. In long-term experiments, both substances were replenished every 3 days. The expression of the ASH2L mutants was induced with Doxycycline (0.1 mg/mL).

### SA-β-galactosidase staining

Staining for β-galactosidase activity was conducted using a citrate-phosphate buffer system (110,111). Cells were seeded at low density to prevent excessive cell-cell contact and let sit to adhere overnight. For the staining procedure, cells were washed twice with 1x PBS and then fixed with 0.5% glutaraldehyde at room temperature for 10 min. Subsequently the cells were washed twice with 37°C prewarmed SA-β-gal washing buffer (400 mM Na2HPO4, 200 mM citric acid, 150 mM NaCl, 2 mM MgCl2, pH 5.5) and incubated with prewarmed SA-β-gal staining solution (8 mL/10 cm dish; 5 mM K-ferrocyanide, 5 mM K-ferricyanide, 1 mg/ml X-gal, in SA-β-gal washing buffer) at 37°C for 20 hours. The cells were then washed with distilled water and pictures were taken under an inverted light microscope at 10x and 20x fold magnitude. Three to five pictures with 200 to 500 cells per sample were taken, randomized and counted by at least three independent people.

### FACS-sorting and pre-staining

For live-cell staining of β-gal activity, cells were pre-treated with 100 nM Bafilomycin A1 for 45 min. Then the cells were stained with C12FDG (66 µM) at 37°C for 45 min and subsequently washed with PBS. For FACS analysis, cells were harvested and washed once with pre-sort buffer (PBS + 3% FCS), pelleted and resuspended at 10^6^ cells per mL in cold pre-sort buffer and filtered through a 45 µm cell strainer. FACS sorting was conducted by the Flow Cytometry Core Facility of the IZKF using the BD FACSAria™ Fusion with the BD FACSDIVA Software Version 9.4 (BD Biosciences, SanJose, CA, USA). The sort was performed at 4°C with a 150 µm nozzle. Cells were detected via FSC, SSC and FITC channel and sorted for high intensity of FSC and C12FDG (FITC). Sorted cells were collected in 1 ml DMEM cultivation medium, pelleted after sorting and resuspended in fresh medium and seeded in 48-well plates for growth analysis in the Incucyte Analysis System.

### Incucyte experiments

Growth curves were recorded in the Incucyte® SX5 Live-Cell Analysis System. Over a time course of two weeks, nine pictures per well were taken by a 10x phase microscope (orange channel; exitation 546 – 568 nm; emission 576 – 639 nm) every two or six hours. Evaluation of cell number and confluency per picture was conducted using the “basic analysis” pipeline of the Incucyte 24A program.

### Cytotox Assays/CellTiter Glo Assays + statistics

To determine sensitivity of dTAG-treated and wild type MEF cells towards cytostatic and senolytic compounds, cytotox assays were conducted with an end-point measurement by using the CellTiter Glo Assay. Cells were treated with Cis-Diammineplatinum(II) dichloride (Cisplatin; Sigma-Aldrich, P4394), Etoposide (LKT Laboratories, E7657), Mitomycin C (Cayman chemical, 11435), Wortmannin (Merck, 681675), Abt-737 (Hycultec, HY-50907) or a combination of Dasatinib and Quercetin (Cayman chemical, 11495 and 10005169). Cells were pre-cultivated in 10 cm dishes for 42 hours with dTAG-13 or EtOH treatment, transferred into opaque 96-well plates at 2000 cells per well with conditioned medium and allowed to adhere at 37°C for 6 hours. Serial dilutions of the different substances were applied in triplicates and incubated at 37°C for 24 hours. Then the cells were washed and fresh medium added containing dTAG-13 or EtOH. Cells were allowed to grow for another 48 hours.

For the analyses, 50 µl CellTiter Glo® reagent was added to each well and plates were shaken at 350 rpm for 10 min to ensure complete lysis of cells. For substances with high staining properties (Abt-737 and Dasatinib), 100 µl of each well was transferred into a new opaque well plate for luminescence measurements. Luminescence signals of each well were measured in a SpectraMax® ID3 Multi-Mode Microplate Reader (Molecular Devices). Mean signals of each treatment was normalized to the mean signals of 6 untreated wells. IC50 values for all compounds were determined in GraphPad Prism and compared using the extra-sum-of squares F test for nonlinear regression models.

### EdU incorporation assay

iMEF cells were treated with or without dTAG-13 and/or Lenalidomide. Three hours prior to harvesting, the cells were incubated with 10 µM EdU (5-ethynyl-2’-deoxyuridine). Cells were then fixed and permeabilized following the manufacturer’s protocol for the Click-iT^TM^ Plus EdU Flow Cytometry Assay Kit (Invitrogen, C10632) and stored in suspension, protected from light, at 4°C. Cell suspensions were further processed according to the manufacturer’s protocol, followed by an additional PI staining for DNA content. Propidium iodide (PI) staining was carried out starting with a 5 min incubation with 20 µg/mL RNase (Invitrogen, REF 8003088) at RT. Subsequently the cells were stained with 50 µg/mL PI (Invitrogen, REF P3566) at RT. Samples were then analyzed by flow cytometry on a BD Canto II flow cytometer.

### Protein analyses

SDS-PAGE and Western blotting have been performed identically to the procedure described previously (21). The antibodies that have been used are listed in Tables 1 and 2. Full size blots are provided in Supplementary Figure S6.

**Table 1:**
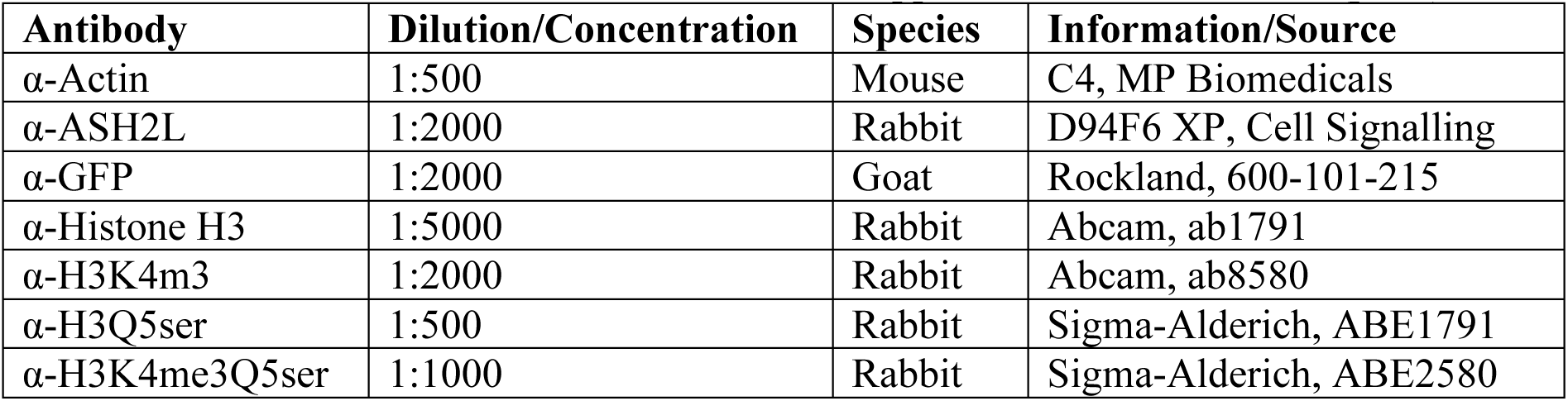
Overview of primary antibodies used for immunoblotting and ChIP experiments. The antibodies were diluted in 1X TBS/T buffer or supplemented with 5% BSA (pH 7).

**Table 2:** Overview of secondary antibodies used for immunoblotting. The antibodies are fused with a horse radish peroxidase (HRP) and were diluted in 1X TBS/T buffer supplemented with 5% nonfat dried milk powder or 5% BSA (pH 7).

| Antibody | Dilution | Species | Information |
| --- | --- | --- | --- |
| α-mouse | 1:10000 | Rat | IgG (H+L) |
| α-rabbit | 1:10000 | Goat | IgG (H+L) |
| α-goat | 1:5000 | Mouse | IgG |

### RNA isolation, cDNA synthesis, and RNA-seq

RNA purification from cells was performed using the HighPure RNA Isolation Kit (Roche, 11826665001). The RNA was transcribed into cDNA by using the QuantiTect Reverse Transcription Kit (Qiagen, 205314). The primers used the RT-qPCR experiments are listed in Table 3.

**Table 3:** RNA-qPCR primer.

| Gene | Primer Name/Sequence | Description/Source (all Qiagen) |
| --- | --- | --- |
| *Rspo2* | Mm_2610028F08 Rik_1_SG | QT00154182 |
| *Angptl4* | Mm_Angptl4_1_SG | QT00139748 |
| *Ccl2* | Mm_Ccl2_1_SG | QT00167832 |
| *Cxcl1* | Mm_Cxcl1_1_SG | QT00115647 |
| *Flrt1* | Mm_Flrt1_1_SG | QT00322266 |
| *Gusb* | Mm_Gusb_1_SG | QT00176715 |
| *Hspa1a* | Mm_Hspa1a_2_SG | QT01660064 |
| *Hsph1* | Mm_Hsph1_1_SG | QT00157367 |
| *Lin7a* | Mm_Lin7a_2_SG | QT01062292 |
| *Mcm6* | Mm_Mcm6_1_SG | QT00102228 |
| *Mga* | Mm_Mga_1_SG | QT01548246 |
| *Mga* | Mm_Mga_2_SG | QT01058806 |
| *Ppbp* | Mm_Ppbp_1_SG | QT00160993 |
| *Tbc1d1* | Mm_Tbc1d1_1_SG | QT00156898 |
| *Vgf* | Mm_Vgf_1_SG | QT00305732 |
| *Vgf* | Mm_Vgf_2_SG | QT01660029 |
| *Wnt16* | Mm_Wnt16_1_SG | QT00134904 |
| *Wnt5b* | Mm_Wnt5b_1_SG | QT00169708 |

Sequencing of 3’-mRNA was performed as described (24). In short, the isolated RNA was analyzed an RNA ScreenTape (Agilent, 5067-5576) with a TapeStation-device (Agilent) for quality control. For internal control, ERCC-RNA-spike-in (ThermoFisher, 4456740) was added to every sample. Libraries were generation using the Collibri 3′-mRNA Prep Kit (ThermoFisher, A38110024). Sequencing was performed on a NextSeq500/550 platform (Illumina) based on a Mid Output Kit 2.5 (Illumina, 20224904) using 75 cycles and single-end reads. Quality control, library preparation, and sequencing were executed by the Genomics Core Facility of the Interdisciplinary Center for Clinical Research (IZKF) of the Medical Faculty, RWTH Aachen University.

### Chromatin immunoprecipitations

ChIP experiments were carried out according to the manual of the ChIP-It High sensitivity Kit (Active Motif, 53040), with a few adaptations. For nuclei isolation, 70 strokes in a 5 mL glass Dounce homogenizer with a tight (B) pestle were applied. Chromatin shearing was conducted using the Bioruptor® Pico sonication device. Sonication was performed in 300 µL aliquots in 1.5 mL Bioruptor® Pico microtube for 4–5 rounds of 10 cycles (each cycle was 30 s sonication/30 s pause) until the majority of chromatin fragments were sheared down to ∼ 200 to 500 bp. Input DNA was precipitated by adding 2 µL of carrier (provided by the kit) and 2 µL glycogen (20 mg/mL). ChIP-qPCR was performed using the QuantiNova SYBR Green PCR Kit. Primers were designed flanking the promotor region of genes using the NCBI Primer design tool. The efficiency of all primers was tested to be above 90%. PCR was run in a RotorGene 6000 (Qiagen). Ct values were determined based on a threshold of 0.01 normalized fluorescence and mean Ct value of duplicate measurement per sample. Expression signal was calculated based on the ΔΔCt method. Signals of immunoprecipitated samples were normalized to signal obtained from input DNA at respective time points. ChIP primers are given in Table 4.

**Table 4:**
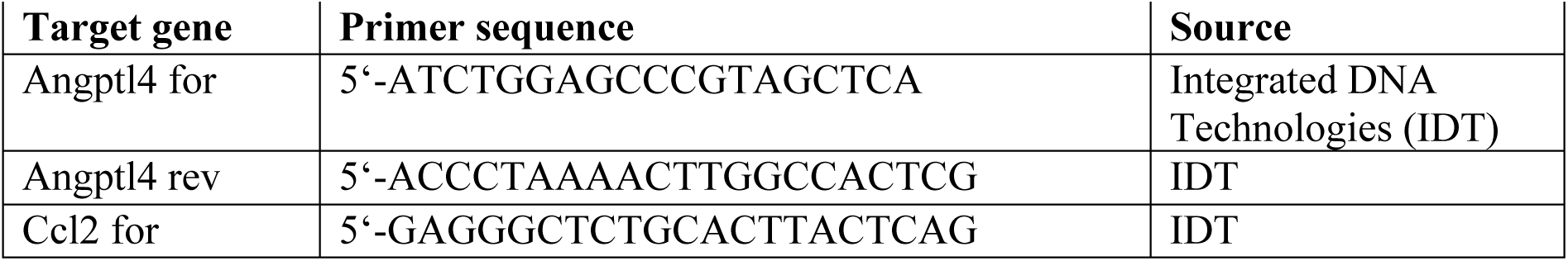
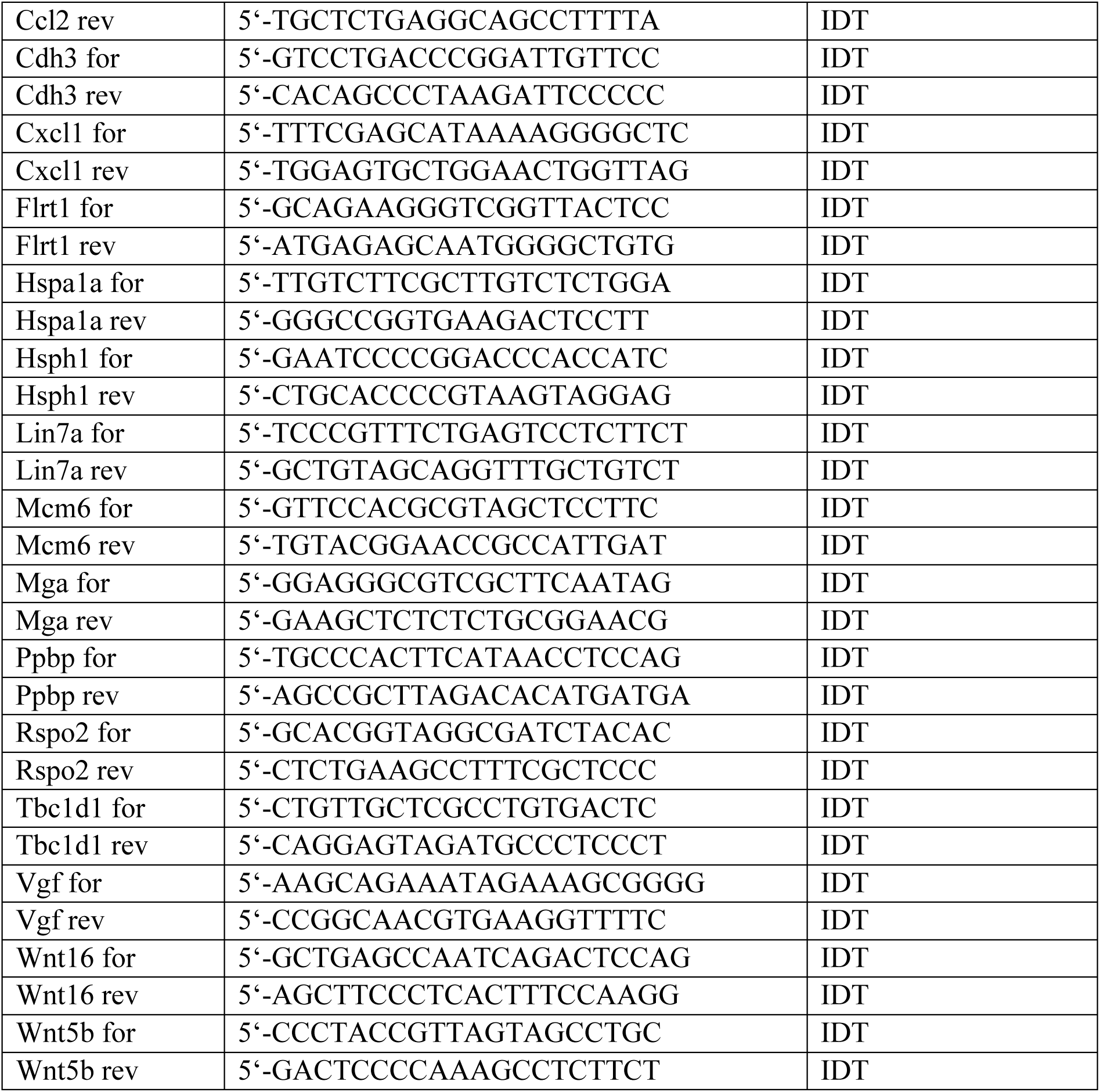
ChIP primer.

### Oligonucleotides

QuantiTect-Primer-Assays were purchased from Qiagen and diluted as indicated in the data sheets provided. Primers ordered from Integrated DNA Technologies (IDT) were diluted to a stock of 100 μΜ in TE buffer (pH 8.0). From these stock solutions, pooled 1:10 dilutions of forward and reverse primer were prepared and employed together. Primers were evaluated for efficiency to be above 95%.

### Single Cell RNA Sequencing and Processing

iMEF cells were treated with or without dTAG-13 and/or Lenalidomide in four biological replicates each as schematically depicted in Figure 4A. The procedure was carried out according to the protocol provided by Parse BioScience (Evervode^TM^ Cell Fixation v3 and Evercode^TM^ WT v2 manual). In brief, cells were fixed, stored at -80°C and subsequently 2000 cells per condition and replicate were further processed. The sequencing libraries were generated with the same Kit in combination with the UDI Plate-WT (Parse BioScience). Quality Control of those libraries was done by TapeStation with the Agilent HS D5000 Screen Tape Assay. Paired-end sequencing of the prepared libraries were carried out using NovaSeq 6000 (Illumina). FASTQ files were generated using bcl_convert (Illumina) and pre-processed using Parse Biosciences split-pipe (v1.5.1) workflow. Reads were aligned using mouse reference genome (GRCm38.p6, GENCODE M20) using STAR Aligner (112). Cell calling and transcript counting were all carried out using the same pipeline. Where applicable, multiple sub-libraries were merged using the split-pipe combine mode.

### Downstream Bioinformatic Analysis

Count matrices were processed using Scanpy (v1.11.5) python package and rapids-singlecell (GPU-accelerated) (113). Quality control was performed per sample, and cells were filtered based on Median Absolute Deviation (MAD). Only cells with a minimum of 500 counts and genes expressed in at least 100 cells were retained. Gene expression was library-size and log-transformed. We identified the top highly variable genes (HVGs) using the ’cell_ranger’ flavor. Principal Component Analysis (PCA) was performed on the HVGs. A k-nearest neighbor (k-NN) graph was constructed, followed by Leiden clustering and UMAP visualization. No batch integration was applied to the data as no strong batch effect was noticed. Cell cycle phases (G1, S, G2/M) were assigned using Scanpy’s implementation of the scoring algorithm based on established marker sets. Senescence-Associated Secretory Phenotype (SASP) activity was quantified using Senepy (v1.0.1). We calculated “Universal” SASP scores to evaluate the senescence burden across treatments (66). Additionally, Decoupler (v2.1.2) was utilized to run AUCell scoring for the “SenMayo” gene set and senescence hubs (63,114).

### Bioinformatics analyses for RNA-seq and ChIP-seq

Data processing: The raw reads of both RNA-seq and ChIP-seq were trimmed using Trim_Galore (https://www.bioinformatics.babraham.ac.uk/projects/trim_galore/). The mouse reference genome mm10 was used (GRCm38.p6; http://ftp.ebi.ac.uk/pub/databases/gencode/Gencode_mouse/release_M25/). The Trimmed reads from RNA-seq were aligned to mm10 using STAR (112). The trimmed reads from ChIP-seq were aligned to mm10 using BWA (115). Samtools were used to generate the sorted Bam files after filtering the unusable reads as recommended by Encode. Quality controls for ChIP-seq were done as recommended by Encode. The replicates were merged using Picard MergeSamFiles (https://github.com/broadinstitute/picard/blob/master/src/main/java/picard/sam/MergeSamFile s.java). Narrow peaks (ChIP-seq H3K4me3) were called using Macs (116). The reads in RNA-seq, which aligned to the mm10 genome, were assigned to the known annotated mm10 genes using featureCounts (117). The counts obtained from featureCounts were normalized using the External RNA Control Consortium (ERCC, Invitrogen) spike-in.

Visualization: DeepTools were used to generate both heatmaps and plotprofiles showing the normalized signal (normalized using CPM (count per Million)) for the ChIP-seq experiment (118). This was centered on TSS of all annotated transcripts in mm10.

The ggmaplot function of ggplot2 package in Rstudio was used to generate the MA plots (119). The apeglm method for log2 fold change shrink age was applied for the MA plots (120). IGV (Integrative Genomics Viewer) was used to visualize the normalized BigWig files to inspect and evaluate the results (121). Screenshot from the Integrative Genomics Viewer (IGV) showing H3K4me3 ChIP-seq signal at representative gene promoters. Signal tracks were generated from BigWig files normalized to 1 million mapped reads. Peaks represent H3K4me3 enrichment at transcription start sites. ChIP-seq depth were normalized to the lowest coverage in each experiment. Differential analysis for RNA-seq and ChIP-seq were carried out using DESeq2 to identify the significantly changed genes or sites, respectively (122). The significantly changed sites from ChIP-seq were then annotated using Homer (123). The information after annotation (distance to the nearest promoter provided by Homer) was used to identify the distance to TSS.

Transcription factor (TF) enrichment analysis: TF enrichment analysis was performed using Enrichr (38,39). The input gene set consisted of 377 genes classified as upregulated without detectable H3K4me3 enrichment at their promoter regions. Enrichment was assessed using the TRANSFAC and JASPAR position weight matrix (PWM) databases, filtered for mouse transcription factors.

For each TF, Enrichr computes enrichment statistics including a Fisher’s exact test p.value and adj.p.value (multiple testing correction), and a combined score, which integrates the significance and magnitude of enrichment. Although several TFs showed nominal enrichment trends, none reached statistical significance based on either p.values. Therefore, TFs were ranked based on both combined score and the number of overlapping genes from the input list. Top ranking TFs ranked were visualized using bar plots.

Gene Ontology analysis: The enrichGO function of the Rstudio’s package clusterProfiler was used to define the enrichd Gene Ontology (GO) terms for biological processes for any interesting group of genes (124). The results were then visualized using the dotplot function of ggplot2 package in Rstudio (119). Parts of the codes used in the mentioned bioinformatics analysis are from nf-core with some modifications (https://nf-co.re/).

## Acknowledgements

We would like to thank J. Vervoorts and P. Korn for helpful discussions, J. Franzen and J. Hübner of the genomics core facility of the Interdisciplinary Center for Clinical Research (IZKF) Aachen, Faculty of Medicine, RWTH Aachen University for sample preparation, sequencing. This work was supported by the Flow Cytometry Facility, a core facility of the Interdisciplinary Center for Clinical Research (IZKF) Aachen, Faculty of Medicine, RWTH Aachen University and by the German Research Foundation DFG, project ID 439895892. We thank N. Steinke of the Flow Cytometry Facility for help with cell sorting. Simulations were performed with computing resources granted by RWTH Aachen University under project rwth0751 (to M.B.). The work was funded by grants from the German Research Foundation DFG (LU466/17-2 to B.L.), the START program of the Faculty of Medicine, RWTH Aachen University (129/22 to M.B.), and institutional core funding (to B.L.).

## Author contributions

B.L. conceptualized the work; J.M., P.B., R.S., A.-M.V., A.T.S., and L.V. performed the wet-lab experiments; M.B., M.H.E.M. and P.K. carried out the bioinformatics analyses; C.K., J.L.-F. and B.L. supervised the work; B.L. wrote the paper; all authors provided descriptions of the Materials and Methods used and commented on the manuscript.

---

**Figure S1 (supporting Figure 1).**
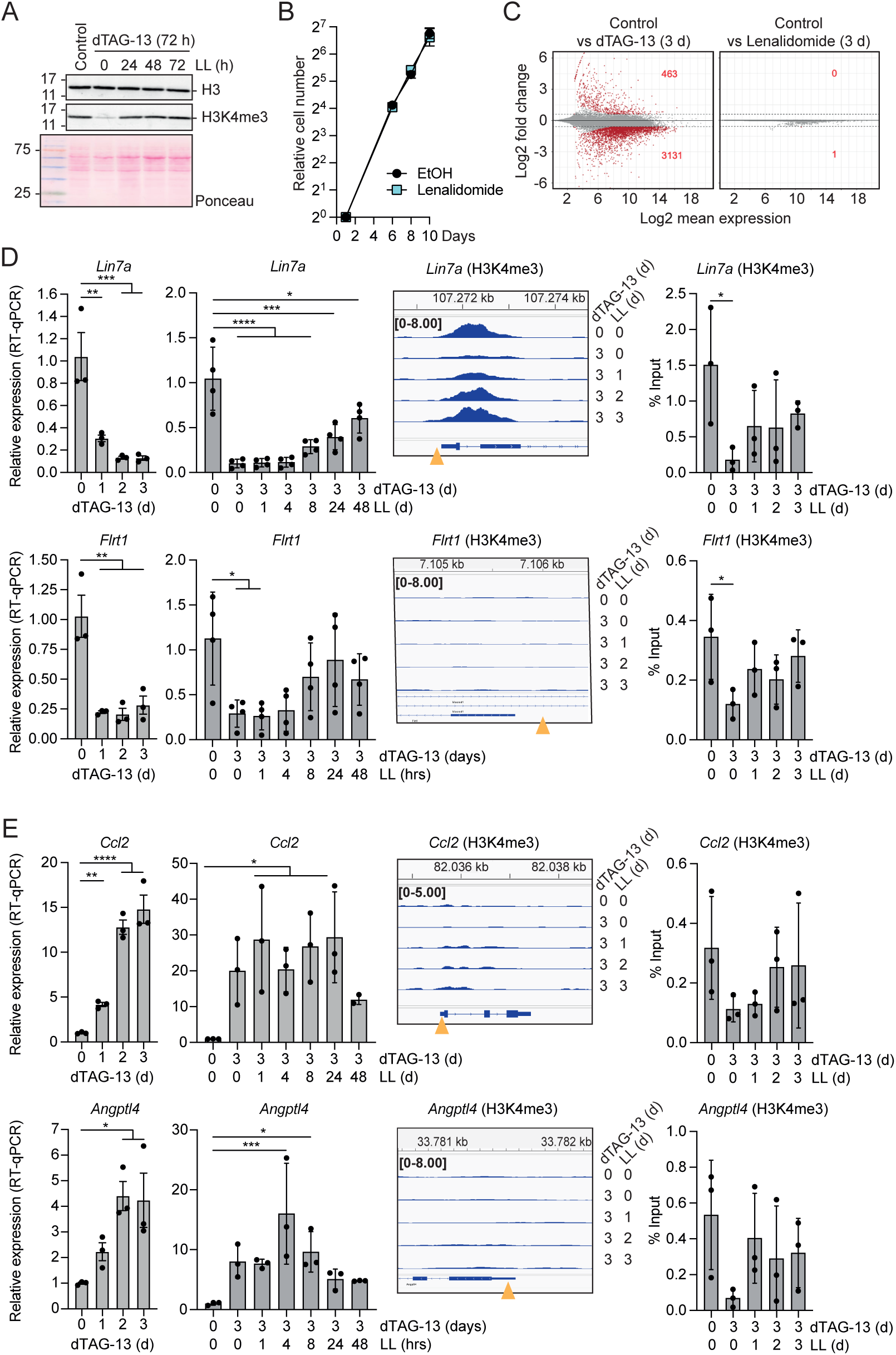
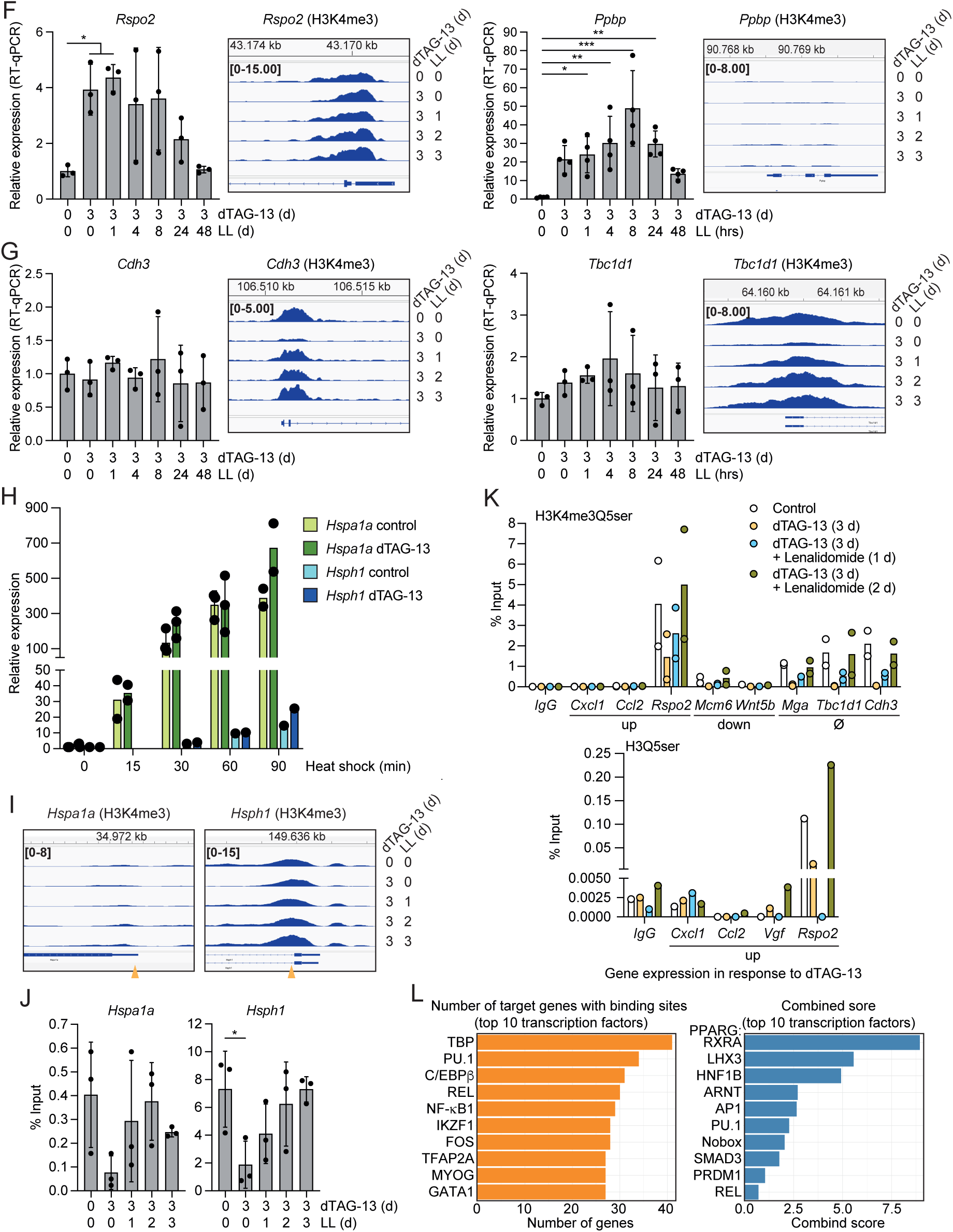
A. iMEF cells were treated with dTAG-13 (100 nM) and Lenalidomide (50 µM) for the indicated times and lysed in RIPA buffer. Histone H3 and H3K4me3 were analyzed by Western blotting. Ponceau staining shows loading control. Uncropped blots are shown in Figure S6. B. Cells were incubated with or without Lenalidomide and counted at the indicated days (3 technical replicates). C. RNA-seq analysis comparing RNA expression of control cells with cells treated with dTAG-13 for 3 days or control cells with cells treated with Lenalidomide for 3 days. Significantly differentially expressed genes are highlighted in red (adj. p-value < 0.05 and logFC > 0.58; two biological replicates, normalized with ERCC spike-in RNA). The total numbers of up- and down-regulated genes are indicated. D.-G. Time course of dTAG-13 and Lenalidomide treatment as indicated. RT-qPCR analyses of the indicated genes are shown (left panels). IGV Snapshot of the ChIP-seq data and for some genes ChIP-qPCR experiments are shown. The ositons of the primer pairs in the promoter region is indicated (yellow arrowhead). Three biological replicates, one-way Anova with multiple comparisons. H. Cells were treated ±dTAG-13 and then incubated at 42°C for the indicated times. RT-qPCR analyses of *Hspa1a* and *Hsph1* expression, 2 or 3 biological replicates. I. IGV Snapshot of the H3K4me3 ChIP-seq data of the promoter regions of *Hspa1a* and *Hsph1*. J. H3K4me3 ChIP-qPCR measurements with primer pairs positioned in the promoter regions as indicated in panel I (yellow arrowheads). K. ChIP-qPCR analyses of promoter regions of the indicated genes with antibodies selective for H3K4me3Q5ser and H3Q5ser. L. Transcription factor enrichment analysis of genes upregulated without detectable H3K4me3 at their promoters. The left panel shows bar plots of the top 10 transcription factors (TFs) ranked by the number of target genes overlapping with the input gene set. The right panel shows the top 10 TFs ranked by Enrichr combined score. *p*-values: *≤0.05, **≤0.01, ***≤0.001, ****≤0.0001.

**Figure S2 (supporting Figure 2).**
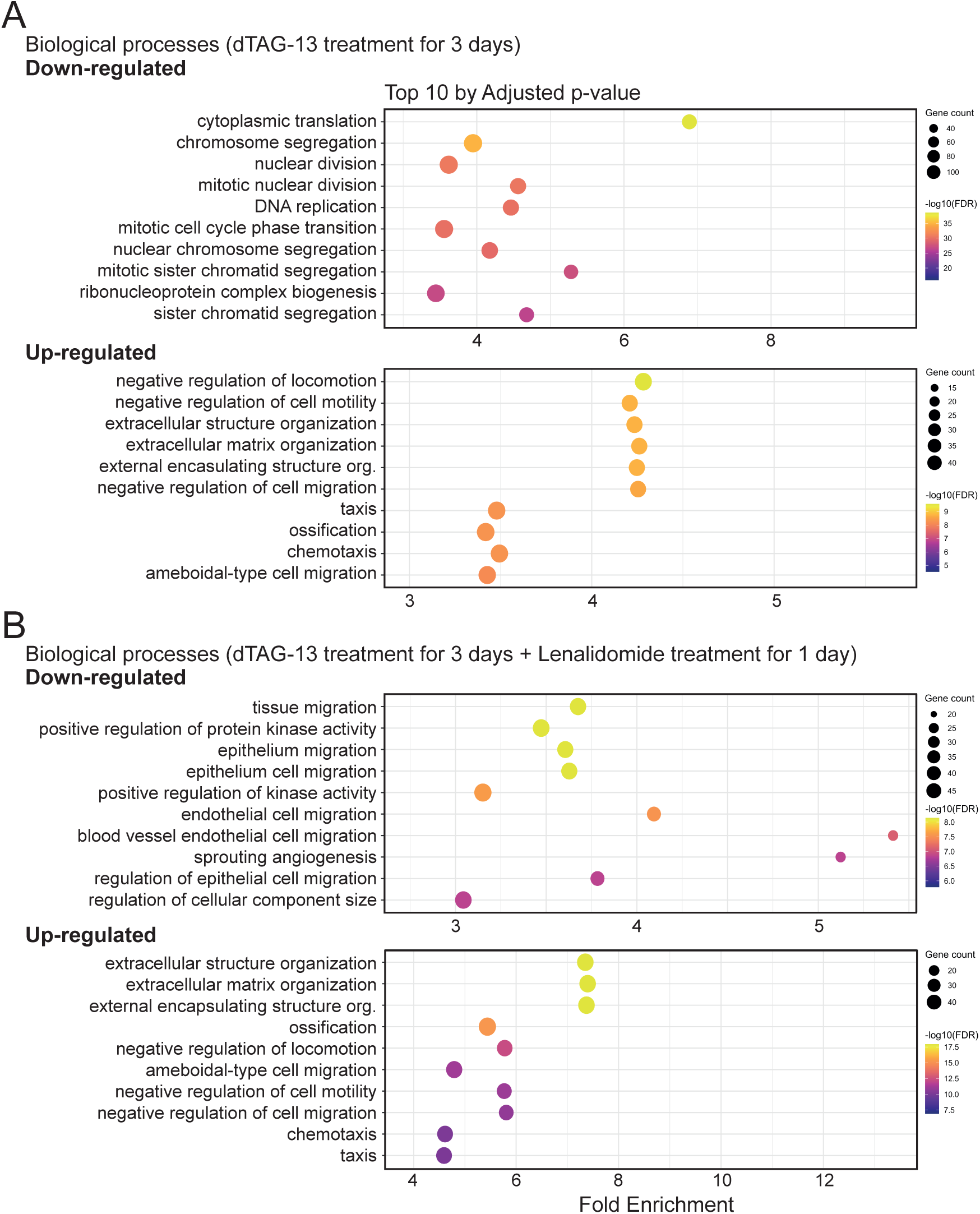
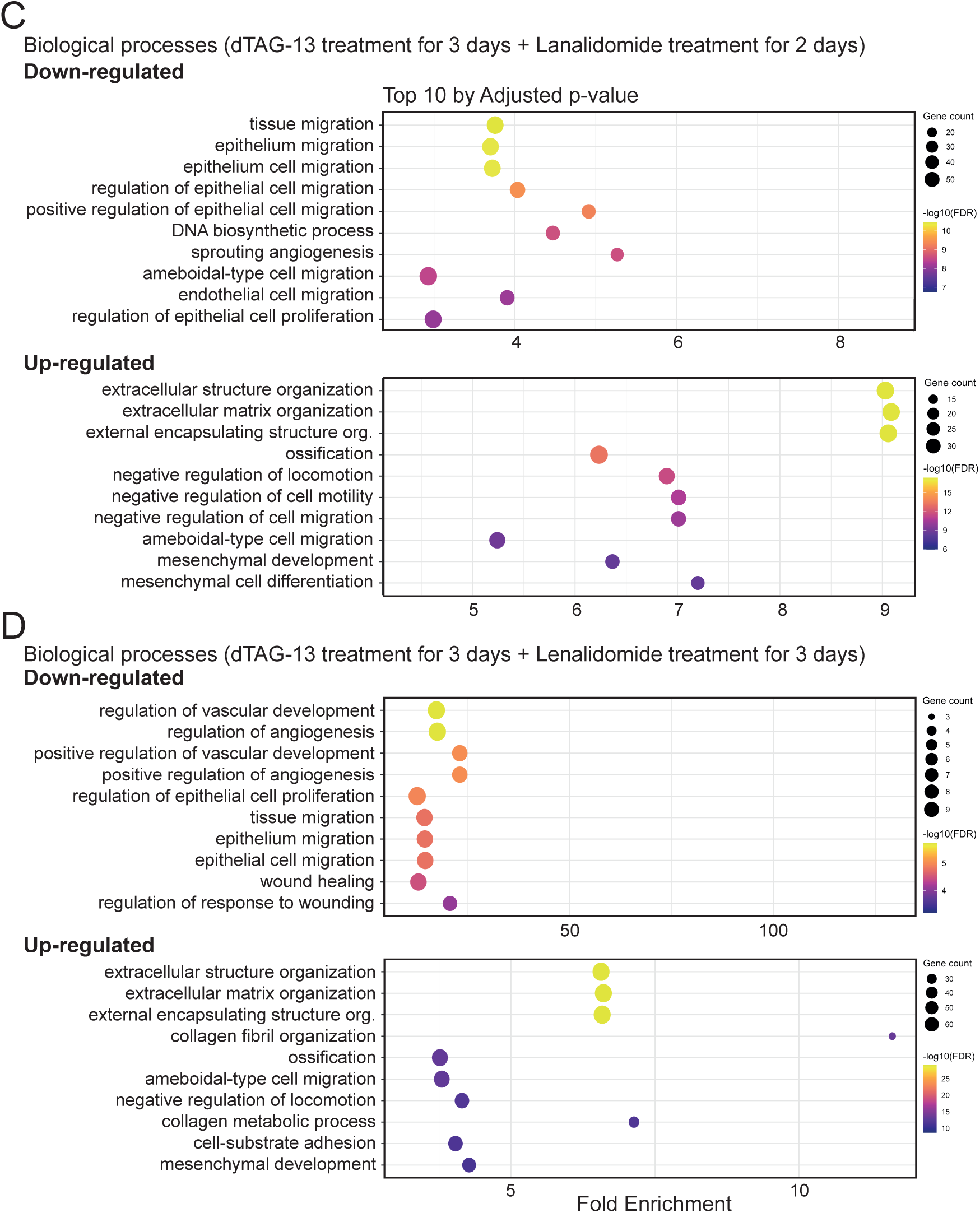
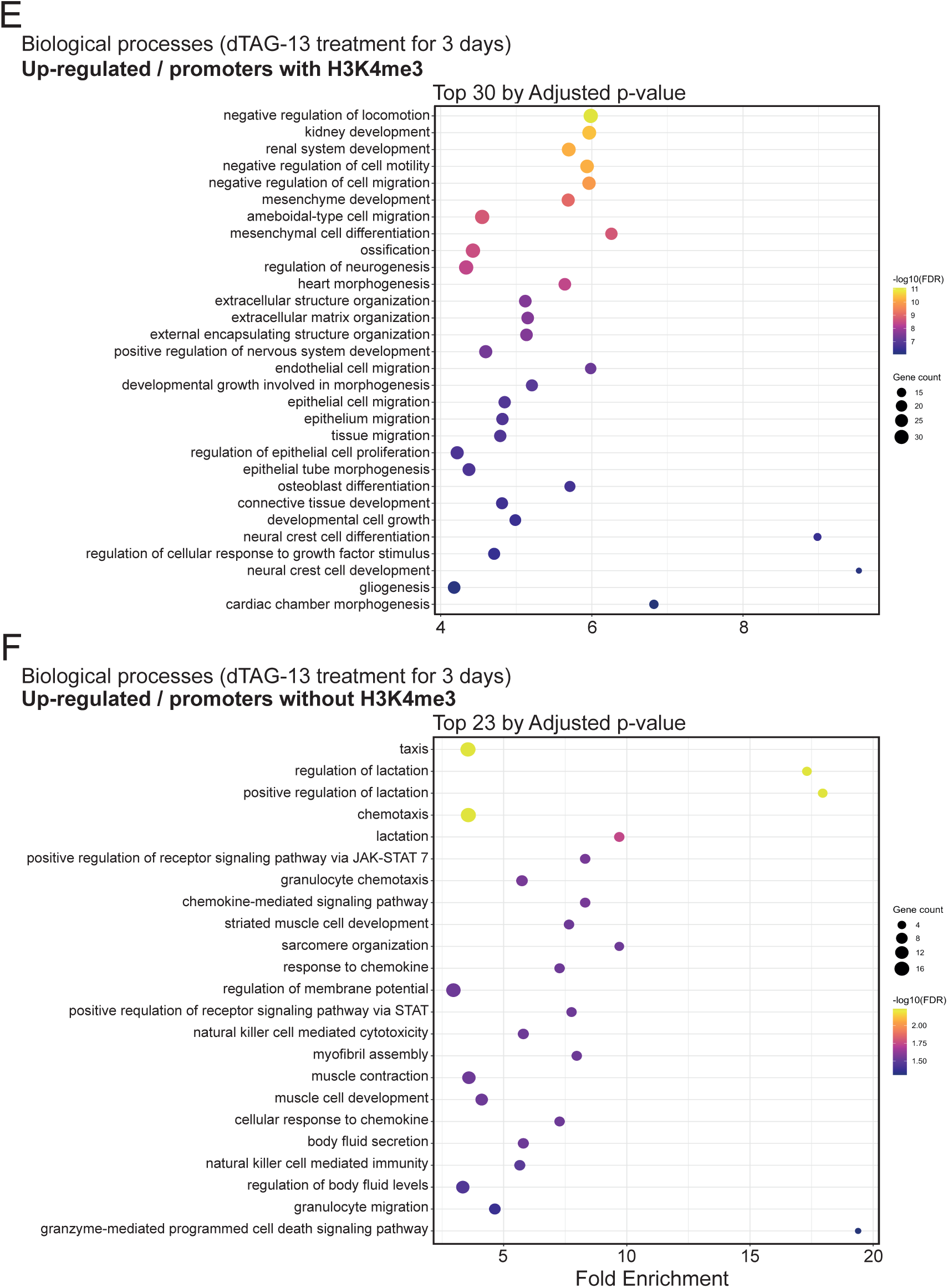
Biological processes in Gene Ontology were evaluated in the RNA-seq data set. Terms were selected based on the lowest adjusted p-value (FDR). Point size indicates the number of genes associated with each term, and point color represents –log10 adjusted p-value. A. The top 10 terms of gene groups of down- and up-regulated genes upon dTAG-13 treatment for 3 days. B. As in panel A, treatment with dTAG-13 for 3 days and subsequently with Lenalidomide for 1 day. C. As in panel B, with Lenalidomide for 2 days. D. As in panel B, with Lenalidomide for 3 days. E. The top 30 terms of gene groups of up-regulated genes with H3K4me3 positive promoters upon dTAG-13 treatment for 3 days. F. The top 23 terms of gene groups of up-regulated genes with H3K4me3 negative promoters upon dTAG-13 treatment for 3 days.

**Figure S3 (supporting Figure 3).**
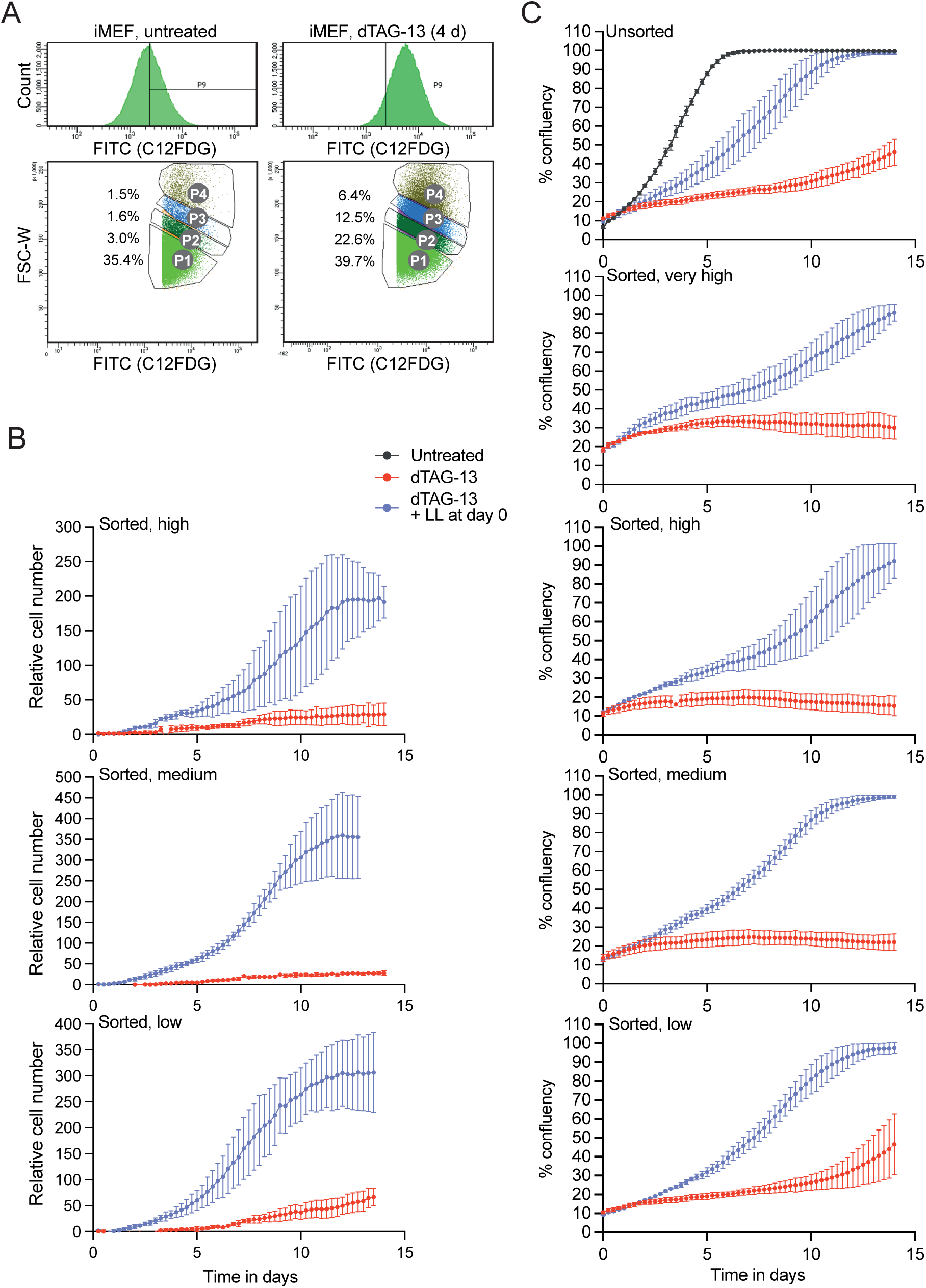
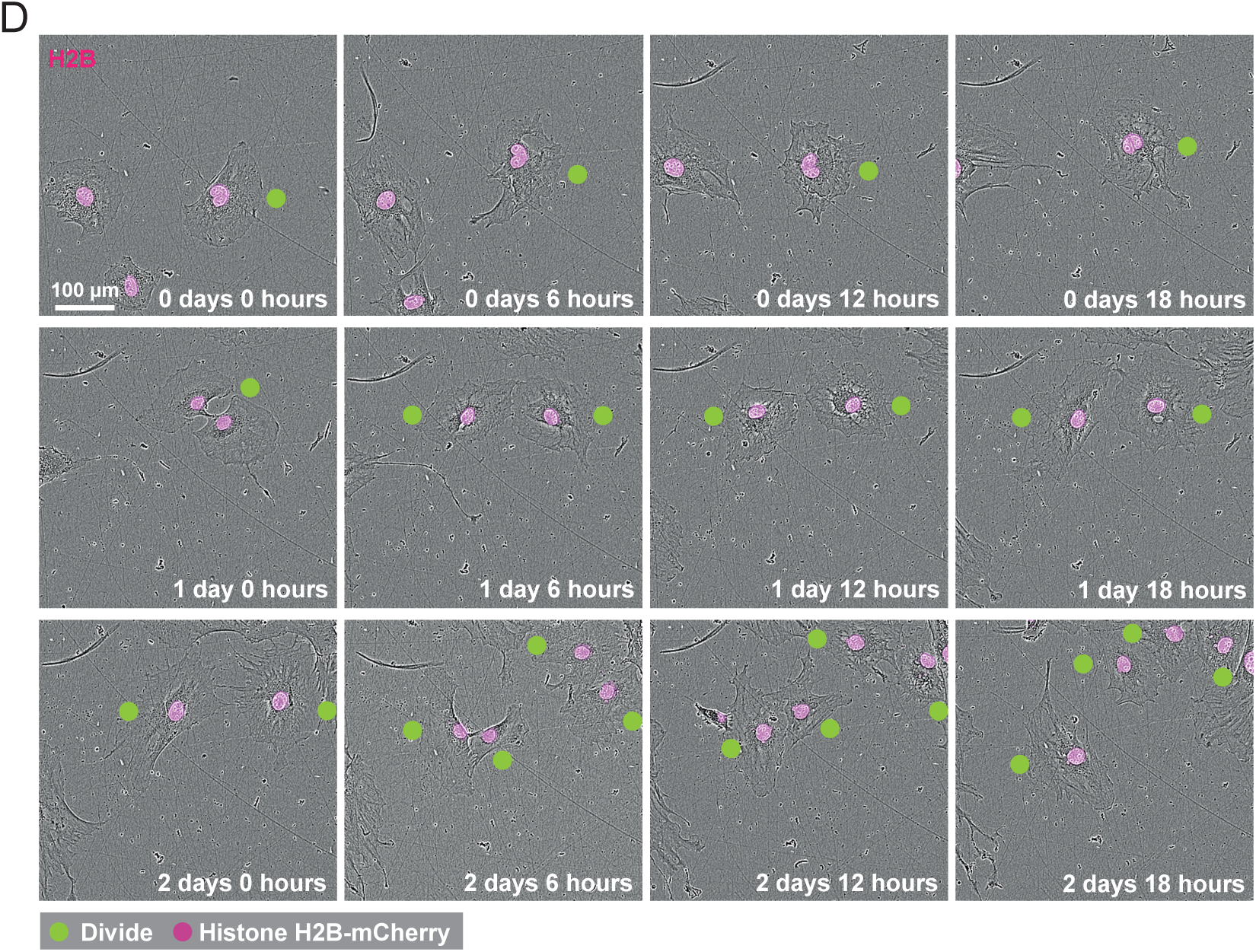
A. Flow cytometry profiles of cells treated ±dTAG-13 for 4 days and stained with C12FDG. The sorted populations P1 – P4 (low, medium, high, and very high group of cells according to SA-β-gal positivity and forward-scatter signals) are indicated. The percentage of cells in the P1 – P4 groups are indicated. B. dTAG-13 treated cells were FACS sorted, replated ±Lenalidomide, and proliferation was documented in the Incucyte SX5 Live-Cell analysis system over 14 days at time intervals of 6 hours. Shown are the growth curves of sorted cells of the low, medium, and high groups. Indicated are mean values ± SD of three replicates. C. As in panel C, but instead of cell number, confluency was analyzed of both unsorted and sorted cells. E. Microscopic pictures of the time course of cells of the very high group. Indicated are cells that divide in the first 66 hrs.

**Figure S4 (supporting Figure 4).**
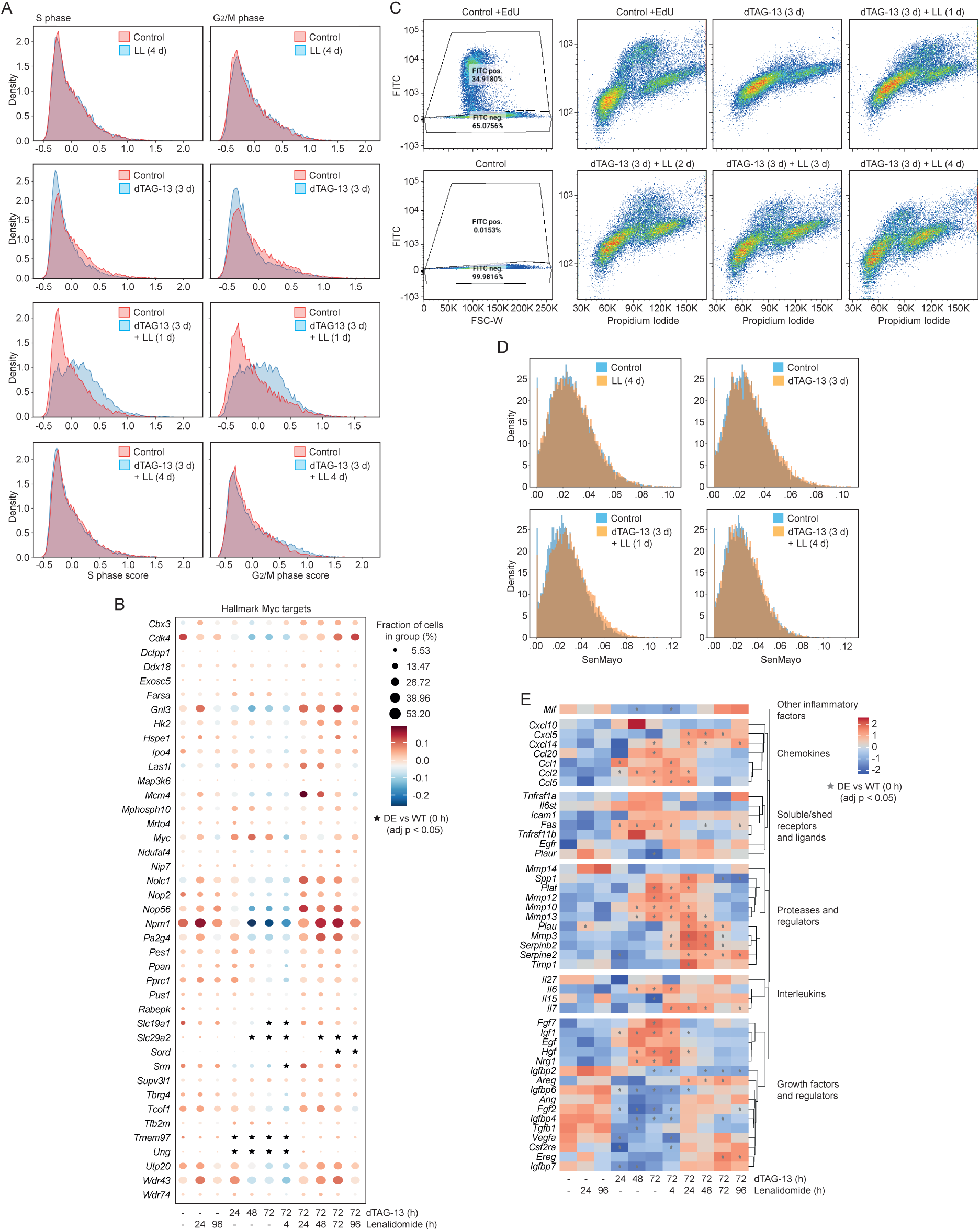
A. S phase scores and G2/M phase scores were analyzed. B. Hallmark Myc targets from the scRNA-seq data set were evaluated. C. Cells were treated with or without dTAG-13 and Lenalidomide as indicated. The cells were labeled for 3 hours with EdU. Subsequently the cells were fixed and EdU incorporation and propidium iodide staining analyzed by flow cytometry. One example of the three biological replicates is shown. The panels on the left show the analysis with and without EdU signals in the FITC channel to determine background labeling. D. The senescence-associated genes in SenMayo (Ref (63)) were evaluated in the scRNA-seq data set of the indicated time points. E. The expression of genes linked to senescence is shown as heatmap.

**Figure S5 (supporting Figure 5).**
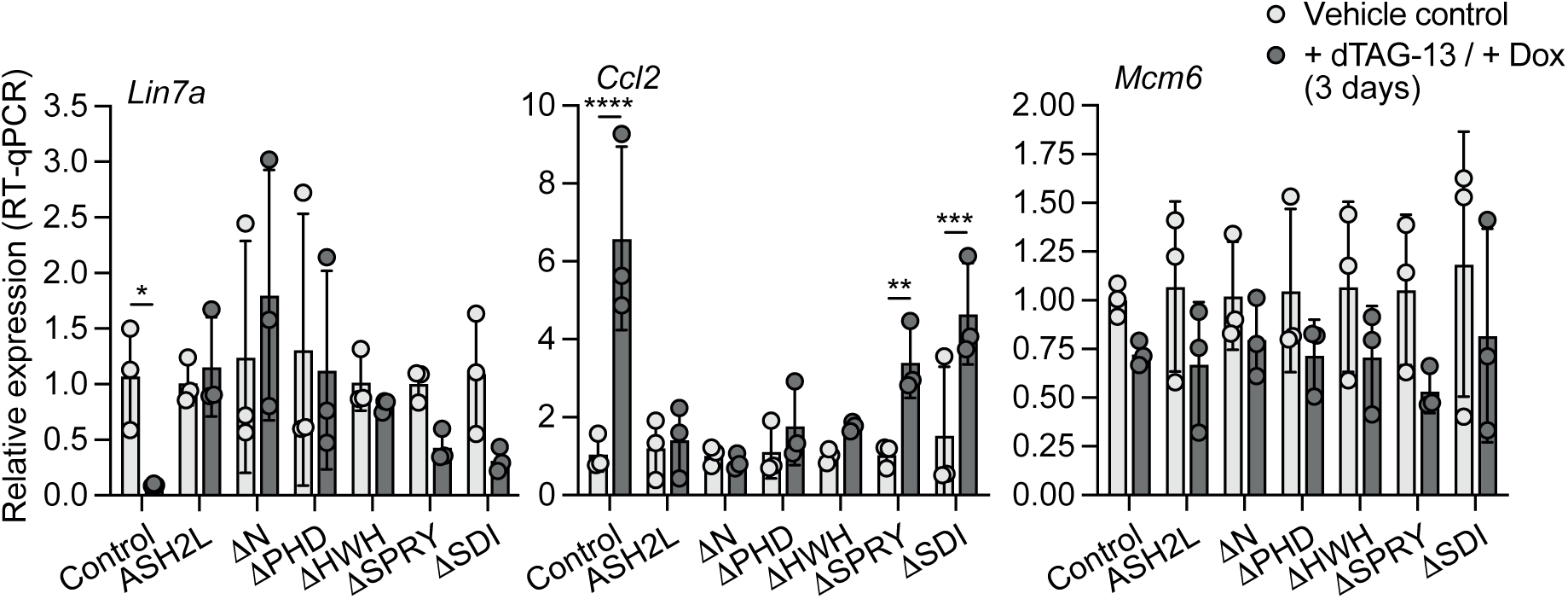
Cells were treated with or without dTAG-13 and Doxycycline for 3 days. The RNA of the indicated genes was analyzed using RT-qPCR. Three biological replicates, one-way Anova/multiple comparisons, mean values ± SD. p-values: *≤0.05, **≤0.01, ***≤0.001, ****≤0.0001.

**Figure S4 (supporting Figures 1 and 5).**
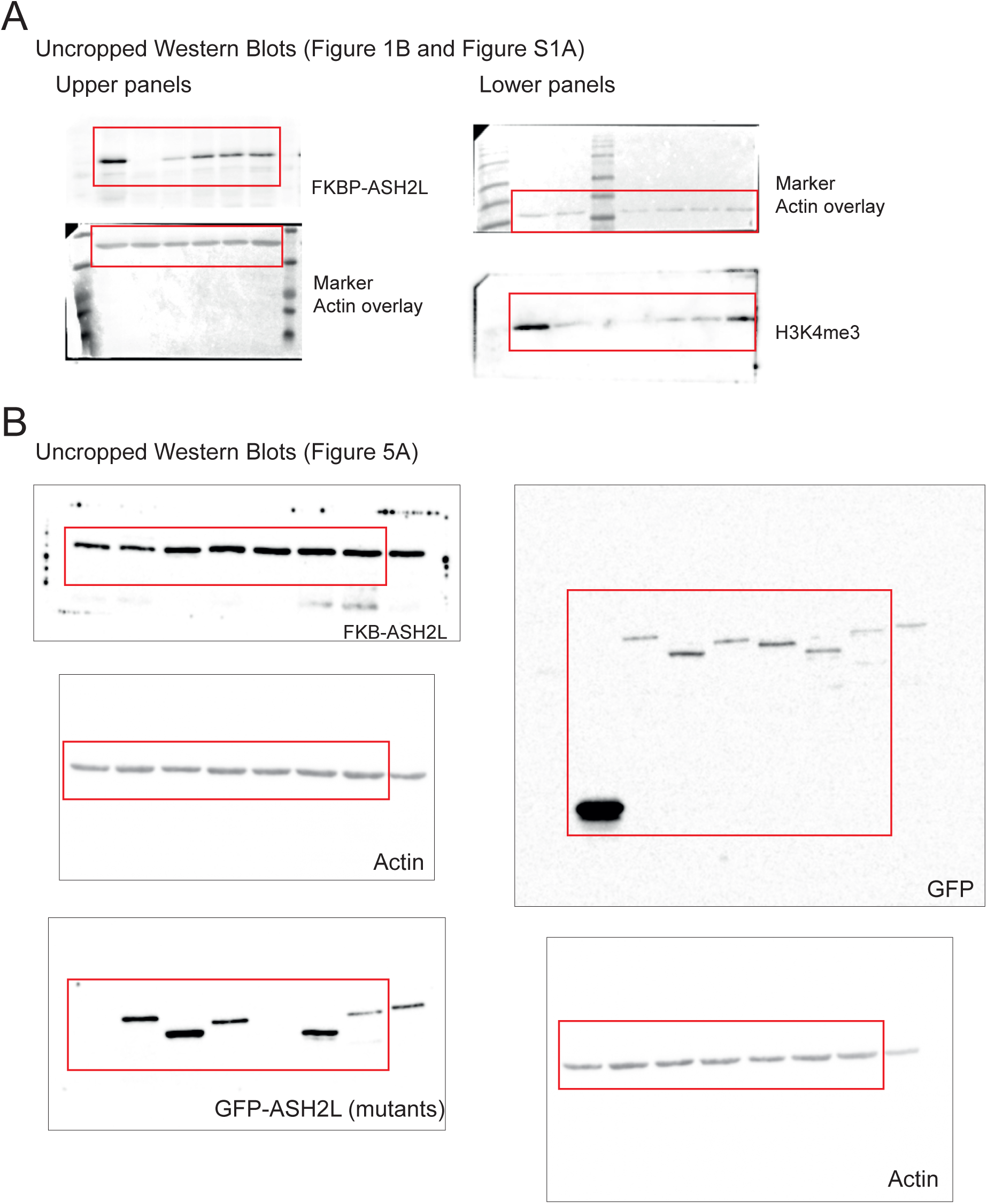

